# Kre28-Spc105 interaction is essential for Spc105 loading at the kinetochore

**DOI:** 10.1101/2021.09.12.459957

**Authors:** Babhrubahan Roy, Janice Sim, Simon J. Y. Han, Ajit P. Joglekar

**Affiliations:** Cell and Developmental Biology, University of Michigan Medical School, Ann Arbor, Michigan, United States of America

## Abstract

Kinetochores are macromolecular protein assemblies that attach sister chromatids to spindle microtubules and mediate accurate chromosome segregation during mit*o*sis. The outer kinetochore consists of the KMN network, a protein super-complex comprising <u>K</u>nl1 (yeast Spc105), <u>M</u>is12 (yeast Mtw1), and <u>N</u>dc80 (yeast Ndc80), which harbors sites for microtubule binding. Within the KMN network, Spc105 acts as an interaction hub of components involved in spindle assembly checkpoint (SAC) signaling. It is known that Spc105 forms a complex with kinetochore component Kre28. However, where Kre28 physically localizes in the budding yeast kinetochore is not clear. The exact function of Kre28 at the kinetochore is also unknown. Here, we investigate how Spc105 and Kre28 interact and how they are organized within bioriented yeast kinetochores using genetics and cell biological experiments. Our microscopy data show that Spc105 and Kre28 localize at the kinetochore with a 1:1 stoichiometry. We also show that the Kre28-Spc105 interaction is important for Spc105 protein turn-over and essential for their mutual recruitment at the kinetochores. We created several truncation mutants of kre28 that affect Spc105 loading at the kinetochores. When over-expressed, these mutants sustain the cell viability, but SAC signaling and kinetochore biorientation are impaired. Therefore, we conclude that Kre28 contributes to chromosome biorientation and high-fidelity segregation at least indirectly by regulating Spc105 localization at the kinetochores.

## Introduction

During eukaryotic cell division, kinetochores facilitate faithful segregation of genetic material from mother to daughter cells. Each kinetochore is a large protein machine that assembles on a specialized chromatin domain called the centromere and establishes end-on attachments between the sister chromatids and spindle microtubules emanating from opposite spindle poles. The budding yeast *S. cerevisiae* has one of the ‘simplest’ kinetochores known to date. Yet, it harbors ∼70 protein subunits. Components of the yeast kinetochore can be divided into two main categories. The first category contains the centromeric DNA binding components and their associated network, known as CCAN (constitutive centromere associated network). The second comprises the microtubule-binding protein network: the KMN supercomplex, the fungi-specific Dam1 complex, and the microtubule plus-end binding protein Stu2 [1–5]. The budding yeast kinetochore incorporates each of these proteins positioned at well-defined average locations along the kinetochore-microtubule attachment (Figure 1A and S1B, [6, 7]). For molecular and cell biologists, the budding yeast kinetochore serves as an excellent model to determine the mechanisms underlying kinetochore functions.

**Figure 1.**
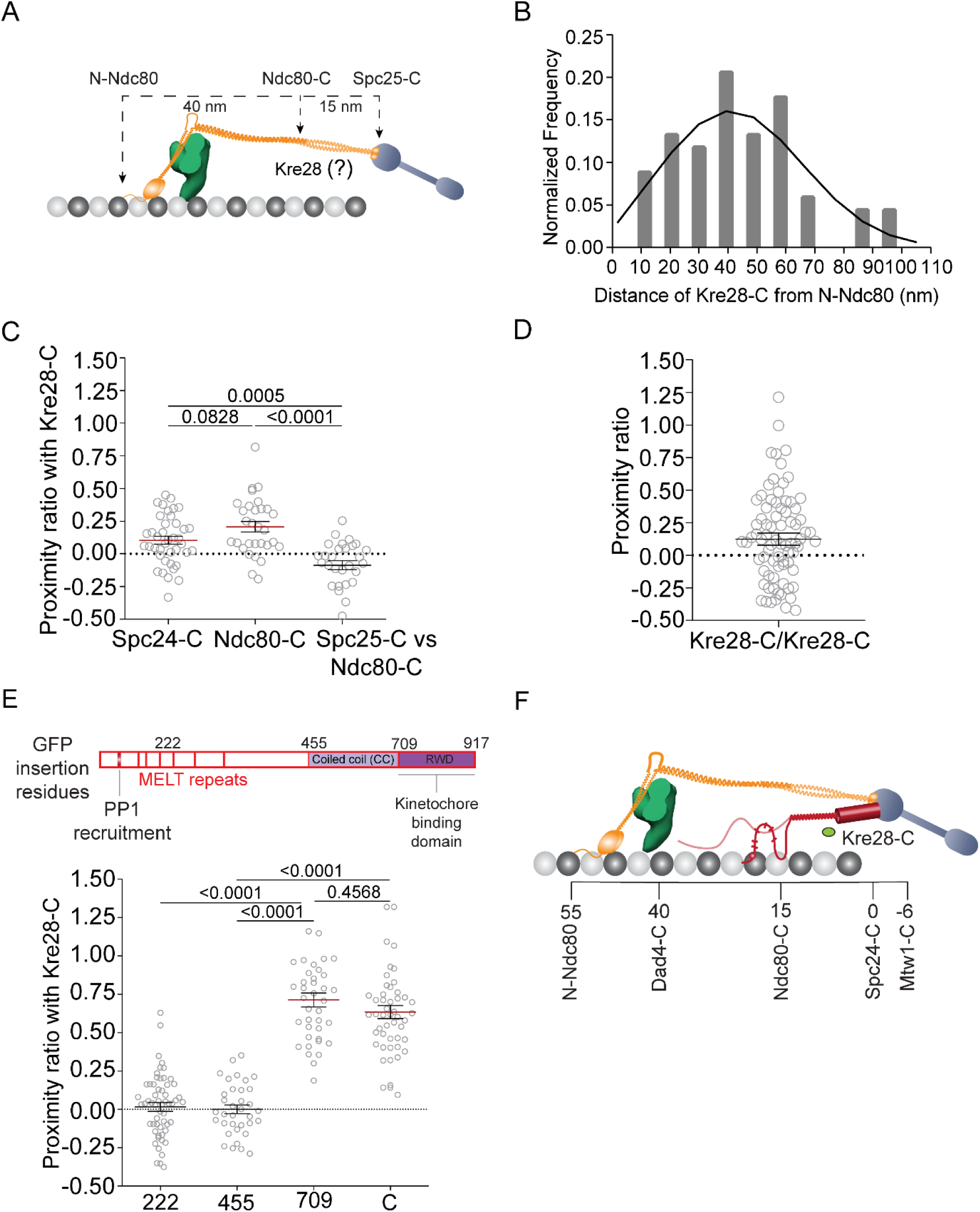
Defining the localization of the C-terminus of Kre28 in the metaphase kinetochore using FRET and high-resolution colocalization. (A) The organization of kinetochore proteins along the microtubule axis in bioriented yeast kinetochores. Positions of C termini of Spc24, Ndc80 and the N terminus of Ndc80 are indicated. (B) Frequency distribution of the distance between the centroids of Kre28-mCherry (Kre28-C) and GFP-Ndc80 (N-Ndc80). The black curve line is the maximum likelihood fit. (C) Proximity ratio for FRET between fluorophores fused to either Spc24-C or Ndc80-C and Kre28-C (mean±s.e.m) in bioriented kinetochore clusters. At least 29 bioriented kinetochores were analyzed for this data set. The p-values obtained from unpaired t-tests are displayed above the plot. (D) Proximity ratio for FRET between adjacent C termini of Kre28 in bioriented yeast kinetochores. 82 kinetochores were analyzed to obtain this data. (E) Top: Line diagram of Spc105 molecule. The illustration was duplicated from our previous study [24]. Red bars represent PP1/Glc7 recruitment site (amino acid 75-79), and six MELT repeats. Amino acid locations of GFP fusion are shown at the top on amino acid residues 222, 455, 709 and C (917). Bottom: Proximity ratio for FRET between Kre28-C and different amino acid positions of Spc105 molecules in bi-oriented kinetochores. At least 35 kinetochore foci were analyzed for this graph. The p-values obtained from one-way ANOVA test performed on the data are mentioned above the plot. (F) Localization of C termini of Kre28 in KMN network of metaphase kinetochores of yeast cells.

Spc105 is an essential kinetochore protein that gets co-purified with COMA complex subunit Mcm21 and with MIND complex (Mtw1-Nsl1-Nnf1-Dsn1) [8, 9]. It forms a complex with another essential kinetochore protein, Kre28, also known as Ydr532C. Kre28 is an orthologue of human Zwint1, *C. elegans* Kbp-5, and *S. pombe* Sos7 [9–12]. Previous studies, using *in-vitro* and *in-vivo* experiments, provide some insights into how Spc105 and Kre28 are assembled at the kinetochores. Still, the specific function of Kre28 remains unclear. How Kre28 localizes within the kinetochore-microtubule attachment site is also unknown [13–15]. Here we define the localization of Kre28 in kinetochore-microtubule attachment sites of bioriented kinetochores and elucidate its functional role.

## Results

### Localization of Kre28 in the KMN network of bioriented kinetochores

We have previously shown that the precise organization and alignment of Spc105 can influence proper SAC activation and silencing [16–19]. Kre28, being an essential component of the kinetochore, may also contribute to the incorporation of Spc105 into the yeast kinetochore. Zwint1, the human orthologue of Kre28, localizes very close to Cdc20 at the human kinetochores [20, 21]. Therefore, Kre28 may play a role in determining the Spc105 organization.

To determine Kre28’s position with respect to Spc105, we first had to define the organization of the entire Spc105 protein within the yeast kinetochore. Previous studies show that the C-terminal RWD domain of Spc105 binds directly to the Mtw1 complex and remains in the proximity of Spc24/Spc25 C-termini [14, 22, 23]. On the other hand, the N-terminus of Spc105 (abbreviated as N-Spc105) consists of a long, disordered phosphodomain that lies somewhere between the Dam1 complex and the C-termini of Ndc80 and Nuf2 within the Ndc80 complex [6, 7, 24]. To map out the overall organization of the Spc105 phosphodomain, we inserted a GFP at locations within Spc105 that demarcate domains predicted to possess secondary structure (see Fig. S1). Additionally, we tagged three different kinetochore subunits to position mCherry at different locations along the kinetochore-microtubule attachment (Fig. S1). Quantification of FRET between the GFP inserted in Spc105 and one of the three mCherry acceptors shows that despite being discorded, the Spc105 phosphodomain localization is mainly limited to a span between the Dam1 complex and the C-terminus of the Ndc80 subunit (abbreviated at Ndc80-C) of the Ndc80 complex. The disordered nature of the phosphodomain also gave rise to FRET between different sections of adjacent Spc105 molecules (Fig. S1D). Having established Spc105, we examined the localization of Kre28 by centroid measurement and FRET assay. Previous literature suggested that the C-terminal structured domains of Spc105 harbor interaction sites for Kre28 [25, 26].

To define the localization of Kre28 within the KMN network of bioriented kinetochores, we performed high-resolution colocalization to measure the mean separation between Kre28-C and the N termini of the Ndc80 subunit (N-Ndc80) in the bioriented kinetochores of yeast (Figure 1A, [7]). We observed that the C-terminus of Kre28 is positioned between 45-50 nm from N-Ndc80, which is consistent with previously published work with Zwint1 [21].

To determine Kre28 localization with higher resolution, we quantified FRET between Kre28-C with either Spc24-C or Ndc80-C in metaphase cells. We obtained a low to moderate proximity ratio in both cases, indicating that Kre28-C may localize somewhere between C-termini of Spc24 and Ndc80 (Figure 1B). The absence of FRET between adjacent Kre28 C termini (Kre28-GFP/Kre28-mCherry) indicated that the C termini of Kre28 molecules are farther apart than 10nm in metaphase (Figure 1C). We also measured FRET between Kre28-mCherry and GFP inserted at different positions of Spc105 (222^nd^, 455^th^, 709^th^, or the C terminus) to find that the C termini of Kre28 are proximal of the kinetochore binding RWD domain (<u>R</u>ING finger, <u>W</u>D repeat, <u>D</u>EAD-like helicases) of Spc105 (Figure 1D and 1E). A previous study suggested that the stoichiometry of Kre28 and Spc105 is 2:1 [11]. However, a comparison of the intensity of Kre28-GFP or Kre28-mCherry signal per kinetochore revealed that there is one molecule of Kre28 per Spc105 molecule in bioriented kinetochores of yeast (Figure S2 A-D).

### Kre28 interacts with the structured middle domain of Spc105 (amino acid 507-638) but not with the kinetochore binding domain

Studies of Zwint1 (orthologue of Kre28 in humans) found that it interacts with a domain within amino acid 1980-2109 of human Spc105 [25]. Protein cross-linking experiments also revealed that coiled-coiled domains Kre28^128–169^ and Kre28^229–259^ interact with Spc105^551-711^, the predicted coiled-coil domain of Spc105 [13]. We wanted to uncover the domains within both Kre28 and Spc105 that are necessary for their mutual interactions.

To study these domains in the ex-vivo condition, we first used the yeast two-hybrid assay. We chose Kre28 fragments (amino acid 1-201 and 202-385, based on predicted secondary structure of Kre28, http://www.compbio.dundee.ac.uk/jpred4/results/jp_OMYEJWN/jp_OMYEJWN.svg.html) and Spc105 <u>c</u>oiled-<u>c</u>oil domain (CC, 455-708) and the C-terminal RWD domain of Spc105 (amino acid 709-917 [14]) (Also see Figure S3A). Both Kre28^FL^ and Kre28^1-201^ showed interactions with CC as indicated by the growth of colonies co-expressing GBD+spc105^CC^, GAD+Kre28^FL^, and GBD+spc105^CC^, GAD+kre28^1-201^ in synthetic dextrose plates lacking histidine. Interestingly, we did not see any interactions between Kre28^FL^ and spc105^CC+RWD^ (Figure 2A). This may be because of the misfolding of spc105^CC+RWD^ fusion with the *GAL4* binding domain (GBD_C1). It is also possible that the RWD domain interferes with the interaction of CC and Kre28, pointing to a regulatory mechanism. To dissect the interaction between spc105^CC^ and kre28^1-201^ more thoroughly, we used smaller fragments (1-126 and 1-80) for our yeast two-hybrid assay with spc105^CC^. We did not notice any interaction using these combinations (Figure 2A). Furthermore, we saw a significant contrast in colony growth between the combinations of spc105^CC^+Kre28^FL^ and that of spc105^CC^+kre28^1-201,^ which denotes a change in the strength of interaction with spc105^CC^. In conclusion, our yeast two-hybrid assay data indicated that Spc105^CC^ binding domain lies within Kre28^127-201^.

**Figure 2.**
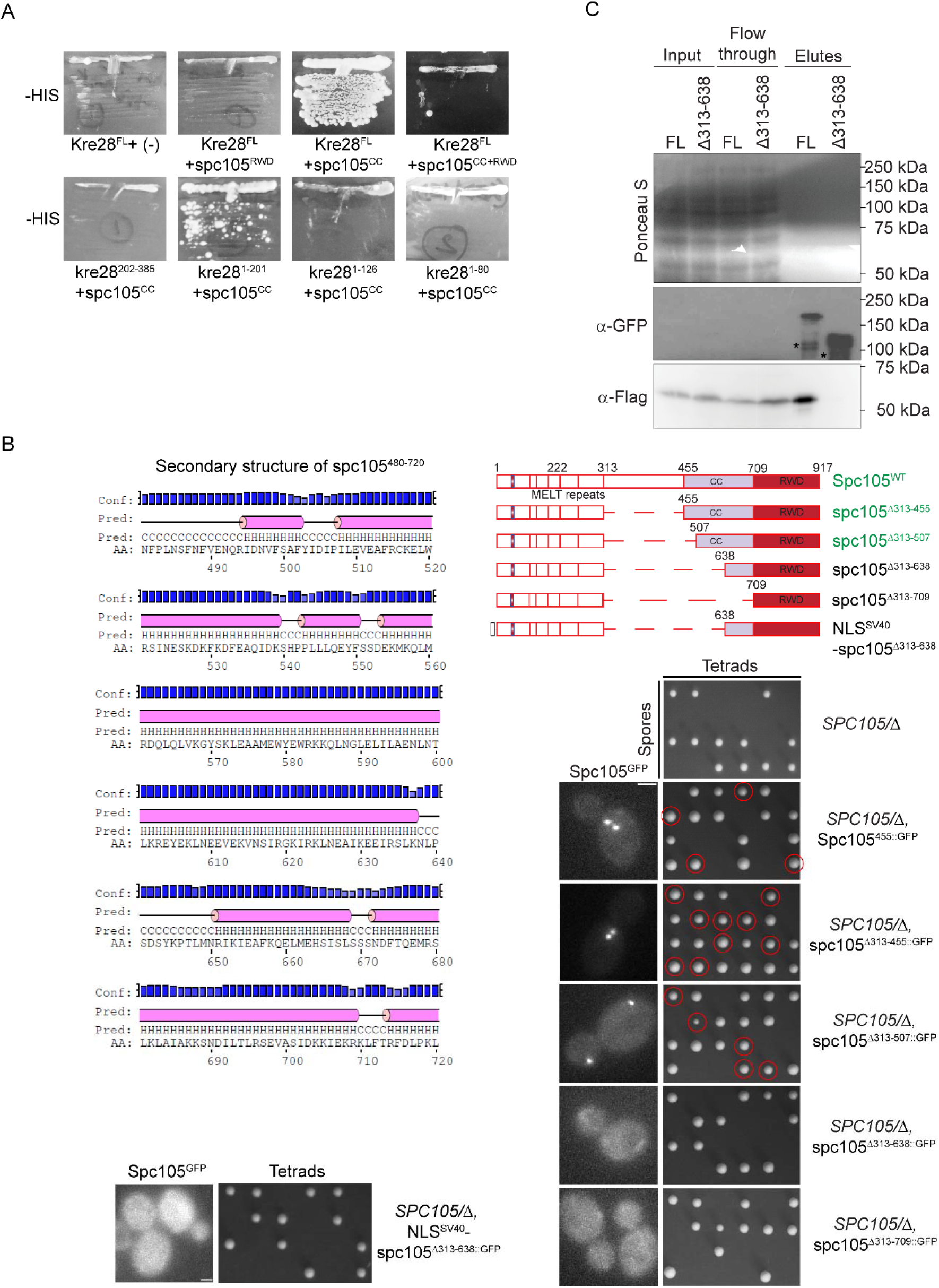
Kre28 N terminus interacts with the helix rich mid strand region of Spc105. (**A**) Yeast two-hybrid assay between Kre28 and <u>coiled-coil</u> domain (CC, Spc105^455-709^) or RWD (kinetochore binding domain, Spc105^709-917^). The plate pictures of synthetic dextrose with histidine dropouts (-HIS) are shown (medium stringent interaction). No growth was observed on plates of adenine dropout (-ADE, high stringent interaction, data not shown). Kre28^FL^ and other kre28 fragments were fused to Gal4 activation domain (GAD, Prey). RWD and CC fragments of Spc105 were fused to Gal4 binding domain (GBD, Bait). Swapping the fragments between GAD and GBD exhibited background growth with kre28^1-80^ fragment on -ADE plates. For controls, please see Figure S2E. (**B**) Top left: spc105^480-720^ harbors CC (spc105^455-709^). a helix rich domain as predicted by http://bioinf.cs.ucl.ac.uk/psipred. Bottom left and right: Domain mapping of Spc105 mid strand unstructured and helical region. Also see Figure S2G. (Top right): Line diagrams of full length Spc105 and the truncations of middle domain. (Bottom right and left): Images of heterozygous diploid strains expressing GFP labelled Spc105 (Wild-type or truncated mutants, genotypes of strains mentioned on the right. For detailed genotype please check Table 2) and tetrad dissection plates of the diploid strains expressing full length or truncated version of Spc105. The genotypes are mentioned on the left of every designated panel. Segregants where genomic *SPC105* is deleted and express truncated version of Spc105 are marked with red circles. (**C**) Interaction analysis of Kre28-5xFlag with Spc105^455::GFP^(FL) or spc105^Δ313-638::GFP^ (Δ313-638) expressed in haploid yeast strains. GFP-Trap assay followed by western blot analysis with anti-GFP and anti-Flag antibodies on the cell lysates, flow through and the elutes of indicated strains. Ponceau S straining of the blot are shown as the loading control. Molecular weights of Spc105^455::GFP^, spc105^Δ313-638::GFP^and Kre28-5xFlag are ∼132 kDa, 93.38 kDa and ∼53 kDa respectively. Due to low expression, we could not detect Spc105 in the Input samples. In the elute samples of both the strains, we observed some bands with lower molecular weight, likely due to the degradation of GFP labeled Spc105.

Next, we mapped the Kre28 interacting domain of Spc105 *in-vivo*. We performed domain mapping experiments where we truncated the mid strand domain of Spc105 (amino acid 313-708) at different residues based on predicted secondary structure (Fig. 2B). We constructed versions of GFP labeled Spc105 with different truncations in the mid strand domain (Δ313-455 harboring only the unstructured region, Δ313-507 containing unstructured region and a small helical domain, Δ313-638 that contains unstructured region and an alpha helix rich domain of CC and Δ313-709 that encompasses the entire mid strand domain, Figure 2B top right) and transformed them in a heterozygous diploid strain (AJY3278, *SPC105/*Δ*::NAT*). We examined the localization of these mutants by microscopy. First, kinetochore localization of Δ313-455 and Δ313-507 displayed no discernable difference in localization compared to wild-type (Figure 2B, right). Our nuclear localization signal (NLS) analyses (nls-mapper.iab.keio.ac.jp/cgi-bin/NLS_Mapper_y.cgi, see the methods section) indicated residues of 337-345 (SSNKRRKLD, score 9.0) and/or that of 599N-625L (score 6.9) contain nuclear localization signals. However, previously we have shown that the mutation of 340-KRRK-343 to alanine residues does not affect the kinetochore localization of the mutant [27]. Therefore, this region of 313-507 is not essential for the kinetochore localization of Spc105. Truncation of 313-638 or 313-709 completely abrogated kinetochore localization of Spc105. According to our analysis, spc105^599-625^ may harbor an NLS, and deletion of this signal may have abrogated kinetochore localization of the mutants expressing spc105 ^Δ313-638::GFP^ or spc105^Δ313-638::GFP^ . We introduced SV40-NLS (NLS^SV40^) at the N-terminus of spc105^Δ313-638::GFP^ to check if nuclear localization of this mutant rescues its loading at the kinetochores (Figure 2B, bottom left). However, we did not observe any kinetochore-specific localization.

Subsequently, we wanted to check whether these truncation mutants can support cell viability. To address this, we induced sporulation/meiosis in heterozygous diploid strains expressing these truncated molecules of spc105 (Δ313-455 or Δ313-507). We observed that they were able to complement the deletion of endogenous *SPC105* (*spc105*Δ). We can conclude that the domain of 313-507 is not essential for any activity of Spc105 that contributes to cell viability. On the contrary, Spc105 mutants with either 313-638 or 313-709 truncated could not rescue the viability of *spc105*Δ. Even fusion of SV40-NLS (NLS^SV40^) at the N-terminus of spc105 ^Δ313-638^ did not rescue its ability to support the cell viability in the absence of wild-type Spc105 (Figure 2B right and bottom left). This data set reveals that the proper localization of Spc105 at the kinetochore is essential for its proper function. They also infer that the lack of nuclear localization cannot explain the lethality of the (spc105^Δ313-638^) mutant.

We confirmed these observations using the plasmid-shuffle assay (data not shown). Therefore, we hypothesized that the domain of Spc105 housed within amino acid 507-638 directly interacts with Kre28. Deletion of this domain abrogates the interaction resulting in delocalization of Spc105 from the kinetochores.

To biochemically confirm the results of the 2-hybrid and localization experiments, we immunoprecipitated GFP-labeled versions of Spc105 from strains expressing either Spc105^455::GFP^(FL) or spc105^Δ313-638::GFP^(Δ 313-638) and examined if both molecules interact with Kre28 (Figure 2C). Immunoprecipitation followed by immunoblot analysis demonstrated that even though Spc105^455::GFP^ binds Kre28-5xFlag, the mutant of spc105^Δ313-638::GFP^ is unable to do so, which indicates that a domain harbored within Spc105^507-638^ is essential for its interaction with Kre28 and subsequently its recruitment at the kinetochores.

### Truncation of Kre28 disrupts its localization from kinetochore. However, truncation mutants of Kre28 support cell viability when over-expressed

Our yeast two-hybrid assay (Figure 2A) indicated that the Spc105 interacting domain of Kre28 lies within amino acid 127-201 of Kre28. The predicted secondary structure of Kre28^FL^ showed that the aforesaid region of Kre28 is helix rich and structured (Figure 3A, http://bioinf.cs.ucl.ac.uk/psipred, also see Figure S3A). To check which domain of Kre28 is essential for its loading at the kinetochore and interaction with Spc105, we created yeast strains that express GFP-fused Kre28 fragments from the *ADH1* promoter. We examined their localization in a diploid yeast strain where one genomic copy of *KRE28* is deleted, and the other allele is tagged with mCherry at its C-terminus (Figure 3B). As expected, GFP-Kre28^FL^ localizes at the kinetochore. On the contrary, we could not detect the localization of GFP-kre28^Δ127-182^or fragments with larger truncations at the kinetochores. When over-expressed from pr*ADH1,* GFP-fused versions of Kre28^FL^ and its truncations revealed a high cytoplasmic GFP signal (Figure 3B). Therefore, we performed similar experiments with the SV40-NLS at the C-terminus of GFP-kre28^Δ127-182^. Even in this case, we did not see any kinetochore localization (data not shown). It should be noted that the GFP tagged Kre28^FL^ and its truncations were expressed at similar levels (Figure 3C).

**Figure 3.**
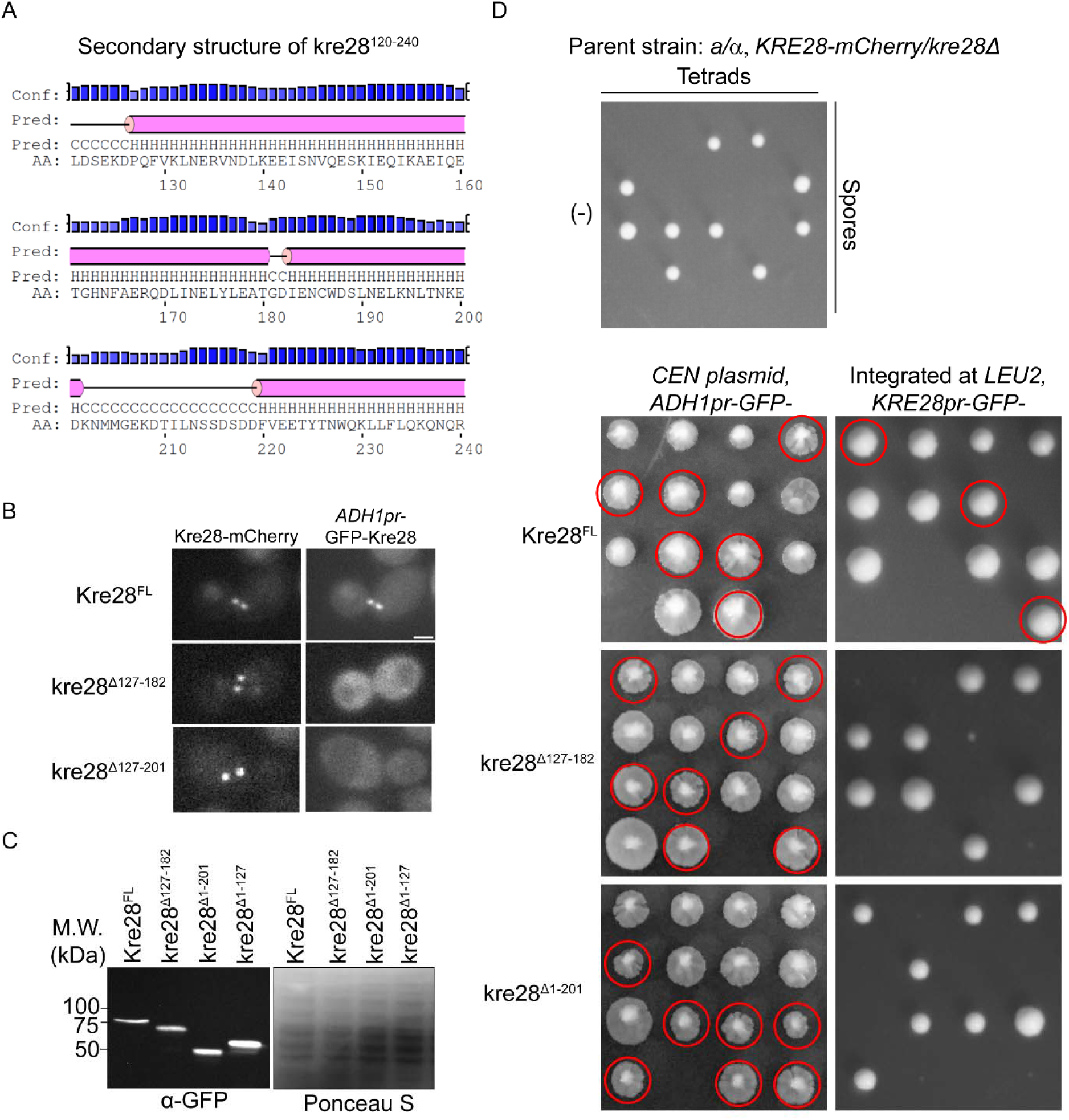
Truncation of Kre28 interferes with its localization to the kinetochore. (**A**) Secondary structure prediction for kre28^127-201^ (http://bioinf.cs.ucl.ac.uk/psipred). (**B**) Representative images of GFP fusions of Kre28^FL^and its truncated versions (kre28^Δ127-183^or kre28^Δ127-201^) exogenously expressed by the *ADH1* promoter. (**C**) Western blot assay with anti-GFP antibody on the lysates of the strains expressing Kre28^FL^ or its truncated version from *ADH1* promoter (*ADH1pr*, over-expression) or its native promoter (*KRE28pr*, expression from *LEU2* locus). Image of Ponceau S stained blot is shown as loading control. Molecular weight of GFP-Kre28^FL^: 73.67 kDa, GFP-Kre28^Δ127-182^: 67.33 kDa, GFP-Kre28^Δ1-201^: 50.6kDa, GFP-Kre28^Δ1-127^: 59.36 kDa. (**D**) Images of tetrad dissection plates for the heterozygous diploid strains. Genotypes are indicated above each photograph. *kre28*Δ spores expressing kre28 truncations are marked with red circles. The plate images on the left were taken after replica plating. Hence, the segregant colonies in those images look larger than the colonies on the right panel.

Does Kre28 delocalization affect the cell viability when yeast cells express the mutants in the absence of wild-type Kre28? To test this, we sporulated these diploid strains and isolated haploid spores. We observed that the segregants over-expressing truncated versions of kre28 rescued the deletion of endogenous *KRE28 (kre28*Δ ). However, the segregants expressing truncated kre28 molecules from their native promoter (*KRE28pr*) could not complement genomic *KRE28* deletion (Figure 3D). These data indicate that full-length Kre28 is essential for binding with Spc105 and its interaction with the Mtw1 complex. However, truncated kre28 mutants with defective kinetochore localization were able to sustain cell viability when overexpressed. We backcrossed viable spores with the parent strain (YEF473) to avoid background mutations. The segregants isolated from those crosses were subjected to further experiments.

### Truncation mutants of Kre28 do not affect kinetochore biorientation but significantly reduce the recruitment of Spc105

We observed slower colony growth among the segregants expressing only the kre28 truncation mutants (Figure 3D and 5B). The slow growth suggested that these mutants have a high propensity of chromosome missegregation, and this will affect their viability. Among these mutants, we chose kre28^Δ127-182^for further analysis because this is the smallest truncation that has a significant defect in kinetochore localization. To check whether kre28^Δ127-182^affects the biorientation of sister kinetochores, we tagged Ndc80 and Spc105 individually with mCherry and studied kinetochore biorientation in kre28 truncation mutant (Figure 4A and 4B). We did not find any noticeable defects in the bipolar attachment of metaphase kinetochores in this mutant (Figure S2F). However, we observed a significant decrease (approximately 61%) in Spc105 recruitment in the truncation mutants (Figure 4B) with a much smaller change (∼ 24%) in Ndc80 localization (Figure 4A).

**Figure 4.**
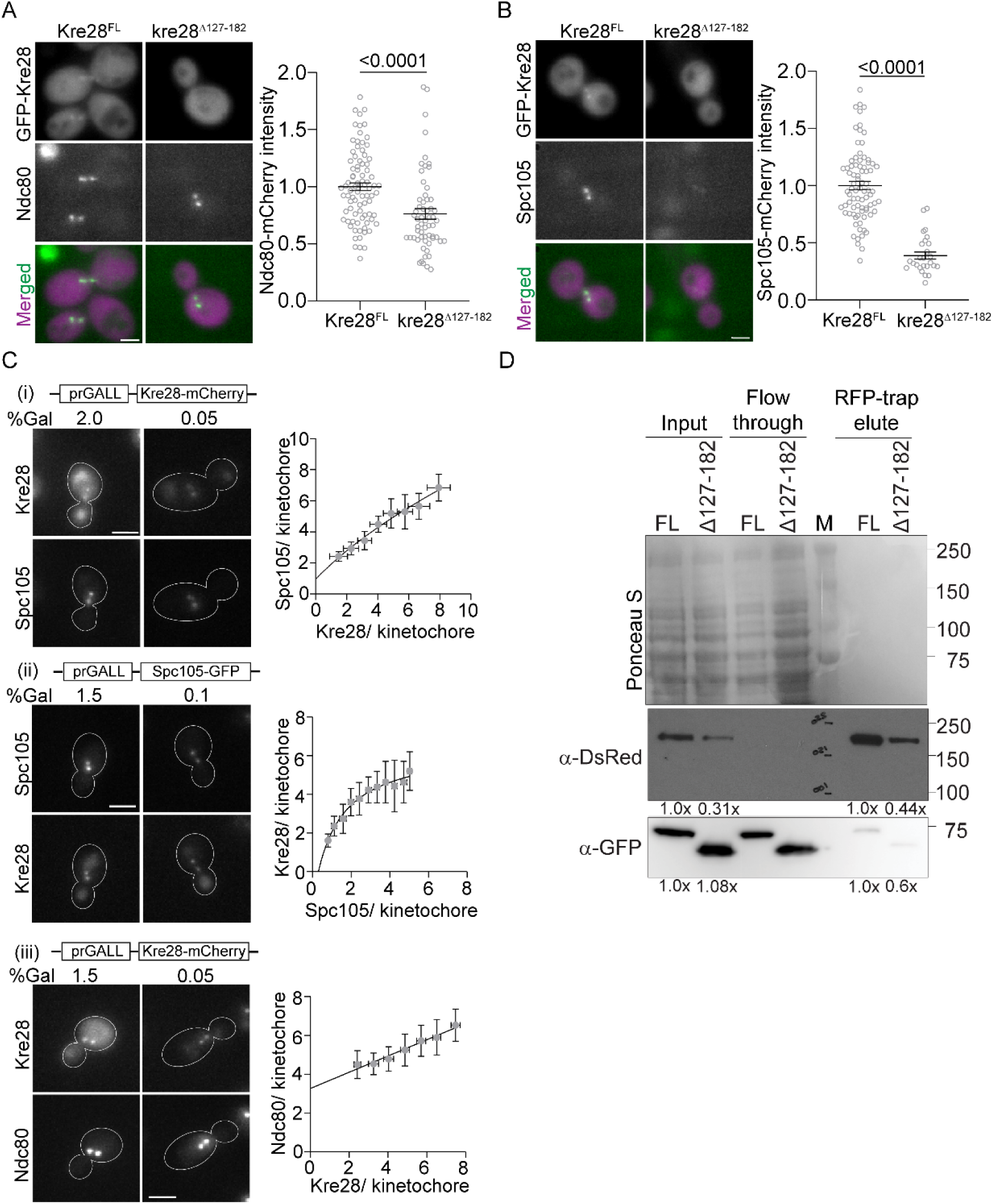
Kinetochores with truncated kre28 mutants form biorientation despite impaired Spc105 recruitment. (**A**) Left: Representative micrographs of GFP-Kre28 (Full length and truncation) and Ndc80-mCherry are shown, scale bar∼3.2μm. Right: Scatter plot of Ndc80-mCherry intensities (mean±s.e.m) is shown for strains with Kre28^FL^and kre28^Δ127-182^. Unpaired t-test revealed p<0.0001, indicated at the top. (**B**) Left: Representative micrographs of GFP-Kre28 (Full length and truncation) and Spc105-mCherry are shown, scale bar∼3.2μm. On the right, scatter plot of Spc105-mCherry intensities (mean±s.e.m) is shown. According to unpaired t-test p<0.0001, indicated at the top. (**C**) (i) Left: Representative micrographs depict Spc105-GFP fluorescence from kinetochore cluster containing different amount of Kre28-mCherry, scale bar∼2.1µm. Right: Scatter plot where each gray circle represents the binned average number of Spc105-GFP molecules plotted against the average number of Kre28-mCherry molecules per bioriented kinetochore. Line in the plot indicates non-linear regression. R^2^=0.9774. (ii) Left: Representative micrographs show Kre28-mCherry fluorescence from kinetochore cluster containing different amount of Spc105-GFP, scale bar∼2.1µm. Right: Scatter plot where each gray circle represents the binned average number of Kre28-mCherry molecules plotted against the average number of Spc105-GFP molecules per bioriented kinetochore. Line in the plot denotes non-linear regression. R^2^=0.9751. (iii) Left: Representative micrographs depict Ndc80-GFP fluorescence from kinetochore cluster containing different amount of Kre28-mCherry, scale bar∼2.1µm. Right: Scatter plot where each gray circle represents the binned average number of Ndc80-GFP molecules plotted against the average number of Kre28-mCherry molecules per bioriented kinetochore. Line in the plot denotes non-linear regression. R^2^=0.9671. (**D**) Immunoblot assay with anti-GFP and anti-DsRed antibodies following RFP-trap assay on the cell lysates, flow through and the elutes of indicated strains. Ponceau S staining of the membrane is shown as the loading control. Molecular weight of Spc105-mCherry, GFP-Kre28 and GFP-kre28^Δ127-182^are ∼132 kDa, ∼74 kDa and 67 kDa respectively. Normalized intensities of Spc105-mCherry and GFP-Kre28 for input and IP samples, which was calculated by ImageJ are depicted below (see Methods).

**Figure 5.**
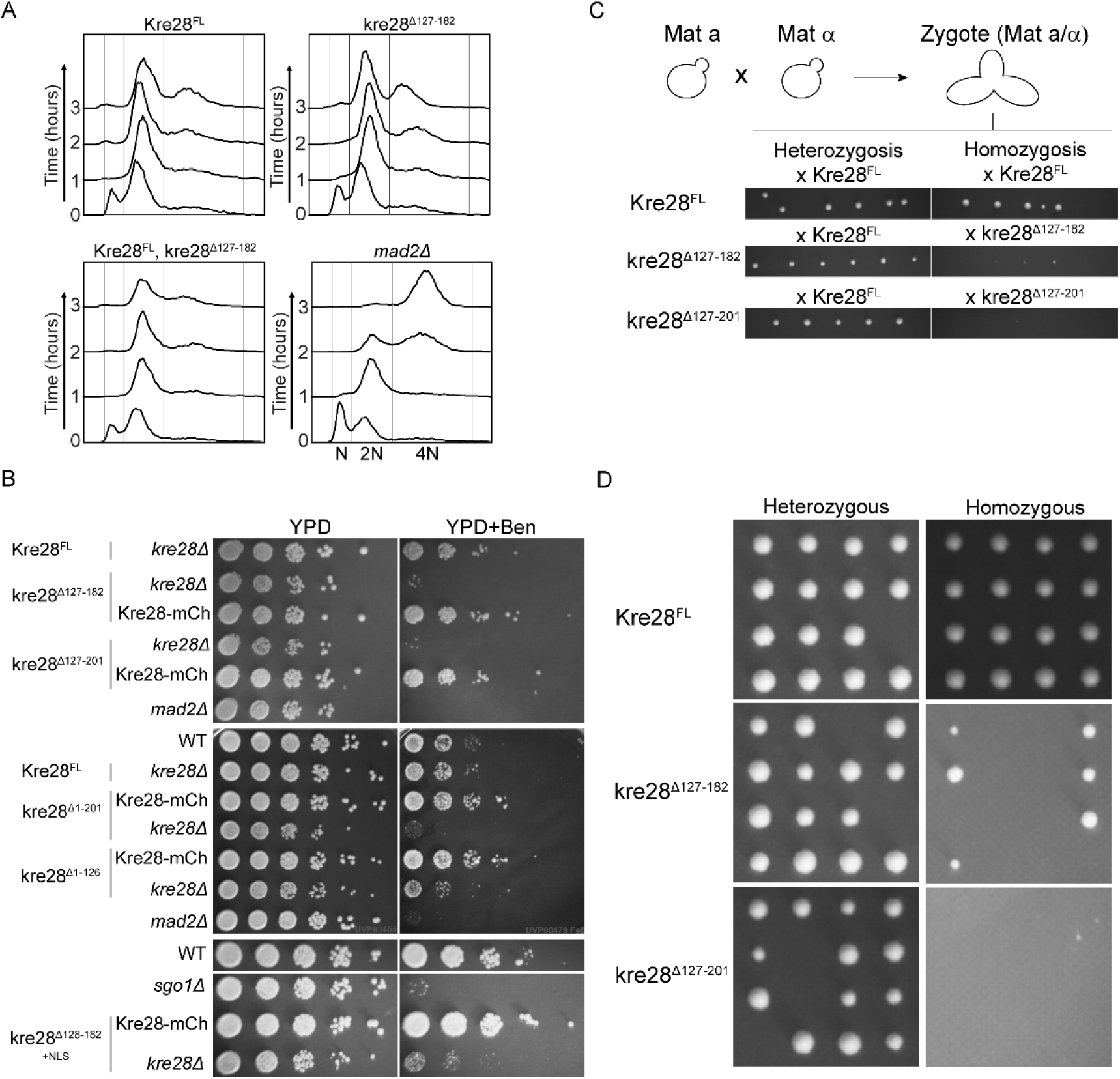
Cells over expressing truncated Kre28 exhibit slower growth and defects in SAC and/or error correction in kinetochore microtubule attachment. (**A**) Flow cytometry showing cell cycle progression of indicated strains that were treated with Nocodazole. The 1n and 2n peaks correspond to G1 and G2/M cell populations respectively. The dominant peak of the 4n in 3h sample of *mad2*Δ strain indicates checkpoint null phenotype. The assay was repeated twice. Presence of a more dominant 2N peak even in untreated samples (0h) of strains expressing Kre28^FL^ may be due to presence of centromeric plasmids, because yeast strains harboring centromeric plasmids typically show a delay in mitosis. (**B**) Benomyl-sensitivity of the indicated strains (see Materials and methods for experimental details. Plates were incubated in 30°C for 2-3 days. The strains of *mad2* Δ and *sgo1* Δ were used as negative controls in these experiments. (**C**) Top: The illustration depicts the zygote formation by mating of haploid strains with opposite mating types which we isolated and incubated in normal non-selective growth media. Bottom: Plate images of crosses between ‘a’ mating type strain of Kre28^FL^ or kre28 truncations and ‘α’ mating type strains with Kre28^FL^ or truncated form of Kre28 are shown. Approximately six zygotes were pulled for each cross. These experiments were replicated three times. (**D**) Plate images of tetrad dissections of homozygous diploid strains (kre28^Δ127-182^ Xkre28^Δ127-182^ ) and heterozygous diploid strains (kre28^Δ127-182^ XKre28^127-182^ ) are shown. To induce meiosis, the zygotes obtained from the crosses were first grown overnight in growth media and then transferred in sporulation media to be incubated for four to five days. After that, tetrads from each sporulation cultures were dissected.

To further characterize the correlation between Kre28 and Spc105 recruitment to the kinetochore, we varied the amount of either Kre28 or Spc105 per the kinetochores by exploiting variable expression of the respective protein using the galactose-induced *GALL* promoter and quantified the amount of Spc105 and Kre28 respectively per bioriented kinetochore (Figure 4Ci and ii). This quantification revealed that the amount of kinetochore localized Kre28 and Spc105 is mutually correlated. As the number of molecules of either protein increases, so does the number of molecules of the other. As expected, both numbers saturate at ∼ 8 molecules per kinetochore, which is close to the maximal number of Ndc80 complex molecules per yeast kinetochore [28]. Given that Spc105 can localize to kinetochores even in strains over-expressing Kre28 fragments, these results strongly suggest that Kre28 positively contributes to Spc105 interactions with the Mtw1 complex. We also found that the number of Ndc80 molecules per kinetochore was slightly lower in cells with kinetochores containing small numbers of Kre28 molecules per kinetochore (Figure 4Ciii). This result is consistent with our prior work, which showed that a reduction in Spc105 molecules per kinetochore similarly lowers the number of Ndc80 molecules per kinetochore [28].

Finally, we performed the immunoblot assay following RFP-trap experiments to assess the severity of impairment in the interaction between kre28^Δ127-182^and Spc105. We observed that the co-precipitation of kre28^Δ127-182^with Spc105-mCherry was significantly reduced (approximately 40%, Figure 4D, please see the figure legend and Materials and Methods section for details of band intensity calculation and normalization). Moreover, we detected the protein level of Spc105 was reduced in the presence of kre28^Δ127-182^(approximately 69%, Figure 4D, panel of anti-DsRed blot). This set of observations imply that the coiled domain Kre28 containing amino acid residue 127-182 plays a significant role in the binding of Kre28 and Spc105. They also suggest that Kre28 plays a role the maintaining the stability of Spc105 protein.

### Truncation of Kre28 adversely affects the spindle assembly checkpoint and error correction pathway and impairs the generation of diploid yeast strains and subsequent meiosis

Spc105 contains short linear interaction motifs known as MELT motifs that, when phosphorylated by the Mps1 kinase, serve as the scaffold for checkpoint components [16, 17]. Moreover, the evolutionarily conserved RVSF motif present in the N-terminus of Spc105 acts as the primary binding motif of protein phosphatase I (PP1) that dephosphorylates the MELT repeats to silence the SAC [19]. Therefore, Spc105 delocalization may affect SAC activation and silencing.

We first studied SAC signaling in cells over-expressing kre28^Δ127-182^by treating cell cultures with the microtubule poison nocodazole and performing flow cytometry to quantify cellular DNA content. The flow cytometry revealed that SAC strength was not discernably different in strains expressing Kre28^FL^and kre28^Δ127-182^, as indicated by the arrest of the cell population with 2N DNA content (Figure 5A). However, we have previously shown that this assay cannot detect smaller defects in SAC signaling [24]. To examine the efficacy to detect smaller defects in SAC and that of the error correction process in kre28 mutants, we subjected them to a low dose of another microtubule poison, benomyl. At its dosage used in this assay, benomyl destabilizes microtubules, and forces yeast cells to rely on a combination of effective error correction and SAC signaling to ensure chromosome biorientation and accurate chromosome segregation[24]. The strains that express kre28 with larger truncations (Δ127-201, Δ1-201, and Δ1-126) demonstrated growth defects even in non-selective growth media (YPD). We also performed the benomyl sensitivity assay using a strain where SV40-NLS is fused to Kre28^127-182^ at the C-terminus. Fusion of NLS also did not alleviate the effect of kre28 truncation, suggesting that the afore-mentioned phenotypes are not caused by delocalization of kre28 from the nucleus (Figure 5B bottom). The growth defects of the kre28 truncation mutants in benomyl-containing media strongly imply that their ability to activate a robust SAC and/or error correction pathway is compromised. Reduced Spc105 loading in the kinetochore of these mutants can explain the observed deficiencies in SAC signaling, error correction, or both.

Previous studies showed that the spindle assembly checkpoint is essential to ensure the fidelity of meiotic segregation [29, 30]. Therefore, delocalization of kre28 may also result in defects in meiotic processes. To examine this aspect, we produced zygotes by crossing *a* and α haploid strains that over-express truncated kre28 in the absence of any wild-type Kre28. However, these zygotes had severe growth defects (Figure 5C). We did not find any nuclear fusion defects during mating of these haploid cells (data not shown). Furthermore, when we sporulated these slow-growing homozygous diploid strains (as shown in one of the panels of homozygosis in Figure 5C) and dissected tetrads, we observed a significant decrease in the number of viable segregants (Figure 5D, right panel). In control experiments, we did not observe any defects in heterozygous diploid formation (Figure 5C, left panel) or the growth of viable segregants after subsequent sporulation (Figure 5D, left panel). We have tabulated our observations for the truncation mutants of Kre28, which we studied here (Table 1). The observation of truncated kre28 mutants affecting meiosis in yeast is similar to what was previously observed in mice oocyte meiosis [31].

**Table 1:** Summary of observations of the experiments involving kre28 truncations.

| Kre28 constructs | Y2H interaction with Spc105<sup>455-709</sup> | Localization at the kinetochore when over-expressed | Sufficiency for cell viability when over-expressed | Localization at the kinetochore when expressed from *KRE28* promoter | Sufficiency for cell viability when expressed from *KRE28* promoter | Sensitivity to Benomyl | Vegetative growth after homozygous mating | Meiosis after homozygous mating |
| --- | --- | --- | --- | --- | --- | --- | --- | --- |
| Full Length (FL) | Yes (*HIS3*) | Yes | Yes | Yes | Yes | No | Normal | Normal |
| Δ202-385 (with NLS) | Yes (*HIS3*) | No | No | Not done | Not done | Technically unfeasible | Technically unfeasible | Technically unfeasible |
| Δ1-201 | No | No | Yes | No | No | Yes | Impaired | Impaired |
| Δ127-182 (with and without NLS) | Not done | No | Yes | No | No | Yes | Impaired | Impaired |
| Δ127-201 | Not done | No | Yes | Not done | Not done | Yes | Impaired | Impaired |
| Δ1-126 | Not done | No | Yes | No | No | Yes | Not done | Not done |

## Discussion

Here we have elucidated how Spc105 and Kre28 are localized and aligned in the microtubule attachment sites of the bioriented kinetochores of budding yeast cells. Kre28-C localizes approximately 48 nm away from the N-terminus of Ndc80. This is consistent with observations reported previously with Zwint1, the human orthologue of Kre28 [21]. Our FRET data are also in accordance with previously published biochemical and cell biological studies [13, 14, 21]. Interestingly, we could not determine the FRET between Kre28-C and components of the Mtw1 complex because strains expressing the fluorescent fusions were inviable. The fluorescent tags most likely disrupt interactions between Spc105/Kre28 and the Mtw1 complex, which results in synthetic lethality [13]. Most strikingly, we observed that the stoichiometry of Kre28 to Spc105 is 1:1, whereas it was previously thought to be 2:1 [11].

Results of the yeast two-hybrid assays involving Kre28^FL^ and Spc105^455-709^ were consistent with previously published data from Yanagida lab [14, 25]. However, it was unclear yeast two-hybrid assay did not work between Kre28^FL^ and spc105^455-917^ (spc105^CC+RWD^). It may be the case that CC+RWD (Spc105^455-917^) fusion with *GAL4* binding domain (GBD_C1) did not fold in a way that they can interact with Kre28. On the other hand, it is also possible that RWD interfering with the binding of CC and Kre28 has an unknown physiological significance. Although we obtained yeast two-hybrid interaction between kre28^1-201^ and spc105^455-709^, we did not detect any kinetochore localization of kre28^1-201^. It was also not sufficient to support cell viability in the absence of wild-type Kre28. Much to our surprise, even after taking the data of Herzog lab into consideration, when we maintained the two binding sites of Spc105 (kre28^128–169^ and kre28^229–259^) in our kre28 cassette (kre28^Δ1-126^) [13], we did not see any kinetochore localization. After considering these data, we can conclude that full-length Kre28 is essential for its localization and for full kinetochore recruitment of Spc105.

Despite being an essential and conserved component of the kinetochore, the structure of Kre28 remains unknown, and the absence of a structure prevents a clear understanding of the implications of our data. Therefore, to provide a structural context to our interaction-mapping and kinetochore localization experiments, we used an implementation of Alphafold2 to predict structures of protein complexes using Google Collaboratory known as Colabfold (https://colab.research.google.com/github/sokrypton/ColabFold/blob/main/beta/AlphaFol d2_advanced.ipynb, [32]). Using Colabfold, we predicted the structure of a heterodimer of Kre28^FL^ and the C-terminal region of Spc105^451-917^ (Figure S3B). The N-terminal boundary of Spc105 used in the structure prediction was chosen based on the start of the structured region within Spc105. Alphafold predicts a ‘T’ shaped architecture for the Kre28-Spc105^451-917^ heterodimer with high confidence for most residues predicted to possess secondary structure (scores 97%-23%, Figure S3B-C). The Pairwise Alignment Error matrix suggests with high confidence that extensive interactions occur among residues in the N- and C-terminal halves of the two proteins. These interactions give rise to two distinct domains within the heterodimer, but the relative organization of these domains cannot be inferred (Figure S3D). The structure correlates very well with the coiled-coil domain predictions for Kre28 and Spc105 (Figures S3A, S3E). It is also consistent with our limited FRET data. We detected significant and similar FRET between Kre28-mCherry and either Spc105^709::GFP^ or Spc105-GFP (Figure 1). In the predicted structure, the separation between Kre28-C and Spc105^709^ and Spc105-C is similar (4.6 nm and 4.8 nm respectively, Figure S3F). Interestingly, we did not detect any FRET between Kre28-mCherry and a Spc105^455::GFP^, even though the model predicts a separation of 7.1 nm between these two points (Figure S3F). However, the model also predicts that the donor and acceptor fluorophores will be in two different domains, and the relative configuration of the two domains is unknown (Figure S3D).

Therefore, the actual separation between Kre28-C and Spc105^455^ is likely to be > 10 nm. The predicted model is also in agreement regarding the residues in Kre28 identified by a previous study to be critical in mediating its interaction with Spc105 [13]. In conclusion, the extensive interaction interface between the two proteins with involvement from residues over nearly the entire length of Kre28 predicted by the model can explain why we could not eliminate the interaction between Spc105 and Kre28 using extensive truncations in either protein.

Our data of Spc105 protein becoming destabilized in the kre28^Δ127-182^mutant (Figure 4D, panel of anti-DsRed blot) implies that impairment of Kre28-Spc105 interaction significantly affects Spc105 integrity of expression. This implication is in accordance with a previously reported study by Zhang and colleagues, where they depleted Zwint1 by RNAi and demonstrated that the cellular protein level of hSpc105 is depleted [33]. A study in the fission yeast *S. pombe* also revealed that kinetochore proteins like Spc105 are surveilled by protein quality control machinery that includes Hsp70, Bag102, the 26S proteasome, Ubc4, and the ubiquitin-ligases Ubr11 and San1 [34]. In that study, the authors suggested that cells employ this mechanism to maintain the homeostasis of nuclear components and genomic integrity. We came across another indirect evidence of protein quality regulation of Spc105 and Kre28 from Yong-Gonzales and colleagues. They showed that both of these proteins become sumoylated by Smc5-Smc6 complex and deleterious mutations in Smc5-Smc6 complex leads to chromosome loss [35]. Consistent with the above-mentioned studies, the Biggins lab also discovered Mub1/Ubr2 ubiquitin ligase complex to be a part of a quality control mechanism that monitors kinetochore protein Dsn1 [36]. A similar mechanism likely controls Spc105 and/or Kre28 levels in *S. cerevisiae*.

Does Kre28 act as a chaperon to stabilize the recruitment of Spc105 at the kinetochore? Some of the previous studies argue against this hypothesis. In human kinetochore, Zwint1 is dispensable for the interaction between hSpc105 and Mis12 complex [14, 15]. While performing ex-vivo kinetochore assembly experiments, Biggins lab showed that Ipl1 phosphorylation of Dsn1, which triggers outer-kinetochore assembly, also recruits Kre28, which should be specific to mitotic cells [37].

Contrastingly, we observed a similar level of Kre28 at the kinetochores at every stage in the cell cycle, including the G1-S phase (unbudded and small budded cells, data not shown), which implies that Kre28 loading at the kinetochore takes place at the same time point as loading of Spc105. Combining our observations with those from Herzog lab, we can conclude that kre28^127-182^ contributes to the main interaction between Kre28 and Spc105 and kre28^229-259^ contributes to interaction with the Mtw1 complex. However, full-length Kre28 is essential for proper binding with Spc105, their mutual recruitment, and their activity at the kinetochores. Kre28 may also become phosphorylated by Ipl1, which can trigger its association with Spc105 and subsequently their loading at the kinetochores.

Does Kre28 have a specific function in spindle assembly checkpoint and error correction or during meiosis? The results of the functional assays (Figure 5) clearly show that the delocalization of Kre28 from the kinetochore impairs the processes. However, all these phenotypes may be linked with the delocalization of Spc105. We also came across the same issue with Zwint1 [21, 33, 38]. Our experiment where we observed a significant number of inviable segregants after sporulation of homozygous diploid strains expressing only truncated kre28 (kre28^Δ^) are also in agreement with a study done by Dong Woo Seo and colleagues [31]. According to that study, Zwint1 depletion results in erroneous chromosome alignment and a high frequency of aneuploidy during mice oocyte meiosis. Our study suggests that this function of Zwint1 is also conserved in yeast Kre28.

## Materials and methods

### Plasmid and Strain construction

The strains and plasmids utilized in this study are documented in table 2 and 3, respectively. Yeast strains containing multiple genetic modifications were constructed by standard yeast genetic techniques. GFP (S65T) and mCherry fusion of proteins were used to localize kinetochores by fluorescence microscopy. The C-terminal tags like GFP, mCherry, 5xFlag and gene deletion cassettes like *spc105*Δ*::NAT* and *kre28* Δ *::NAT* were introduced at the endogenous locus through homologous recombination of PCR amplicons [39]. A 7-amino-acid linker (sequence: ’RIPGLIN’) bridges the tags (GFP, mCherry, or 5xFlag) from the C-termini of the tagged proteins. Earlier, we observed that the intensity of mCherry-tagged kinetochore proteins varies significantly from one transformant to another for the same strain, due to inherent variability of the mCherry brightness. Therefore, to construct all the FRET strains with Ndc80, Stu2, Nsl1, Kre28 and Ask1-mCherry, we crossed a specific mCherry strain with haploid strains of all GFP fused Spc105 alleles and sporulated the heterozygous diploids to obtain the desired segregants.

**Table 2:**
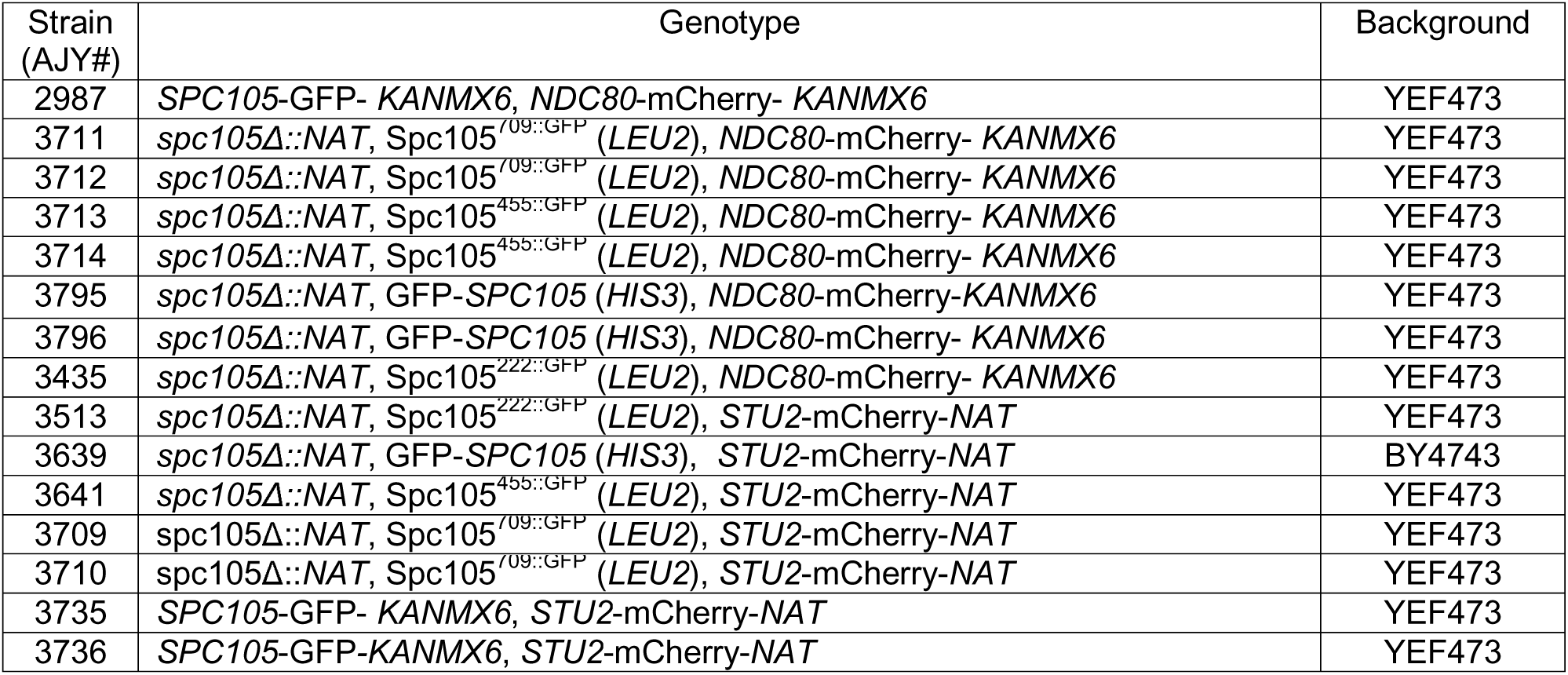
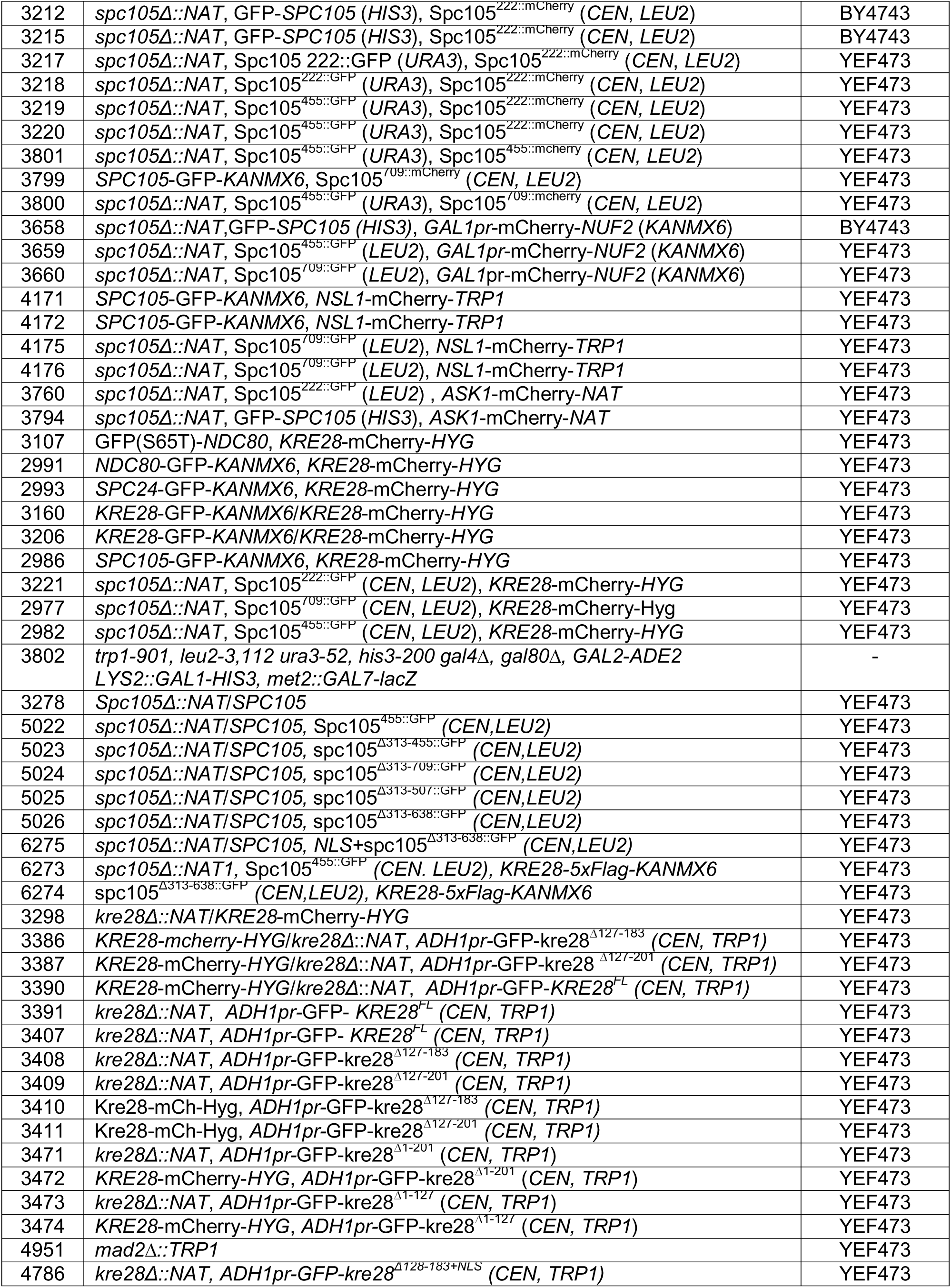
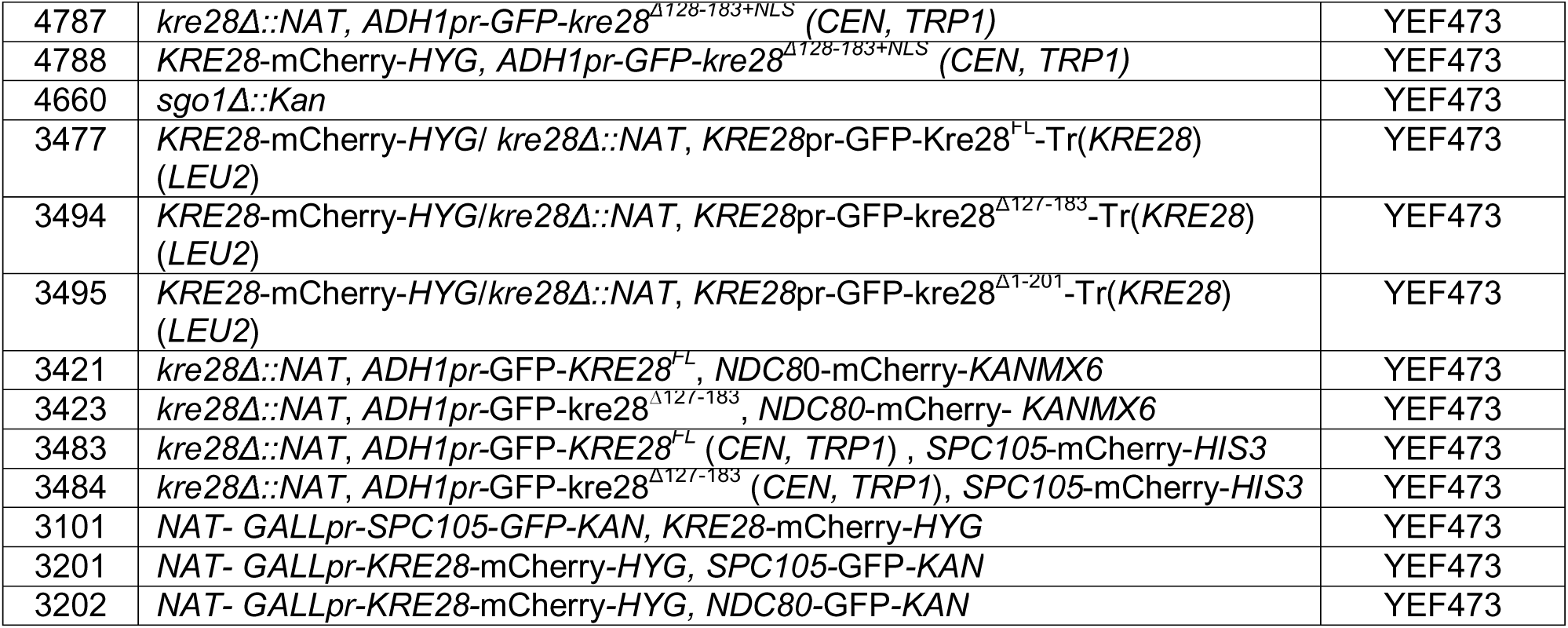
Strain used in this study

**Table 3:**
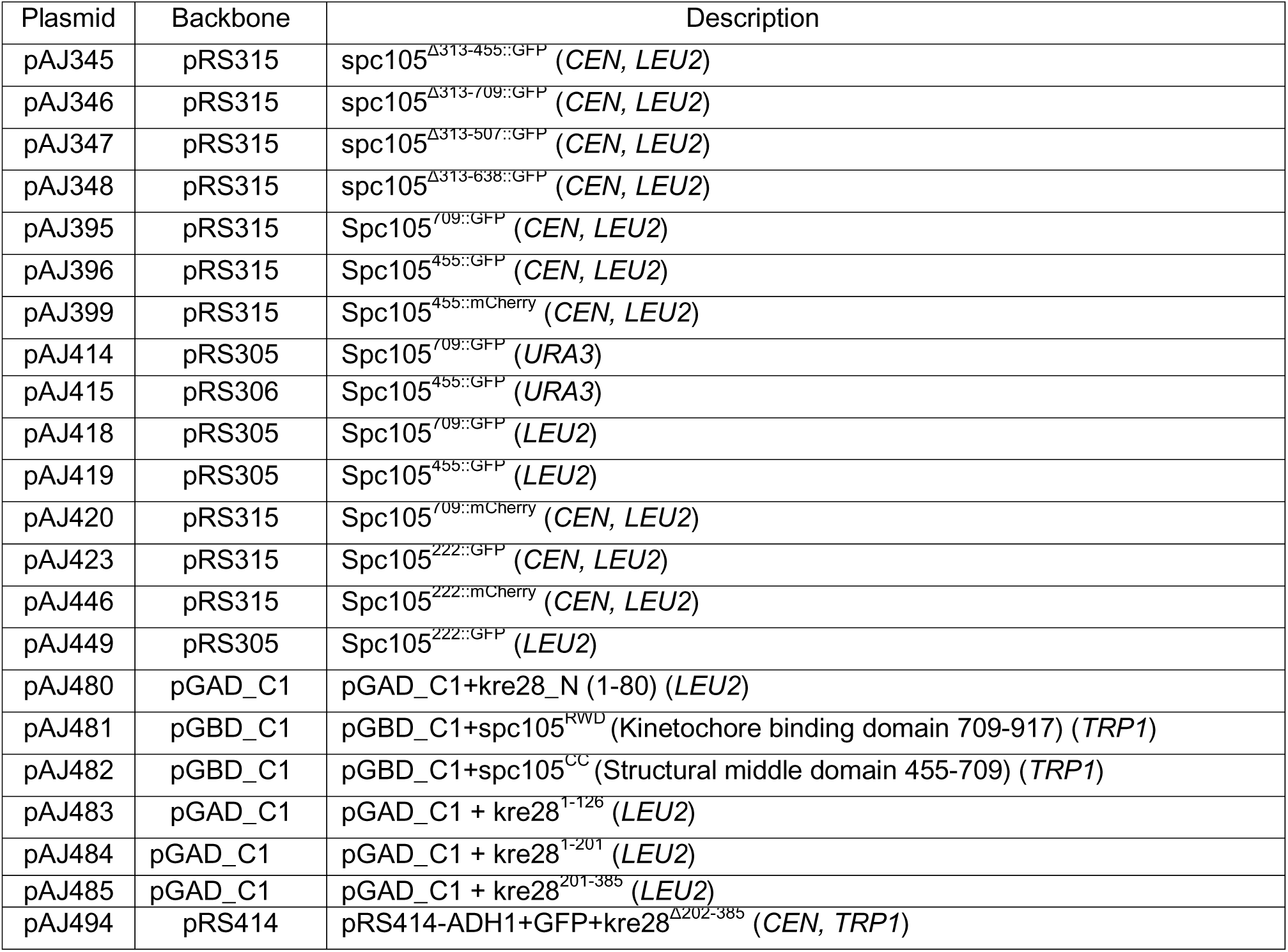
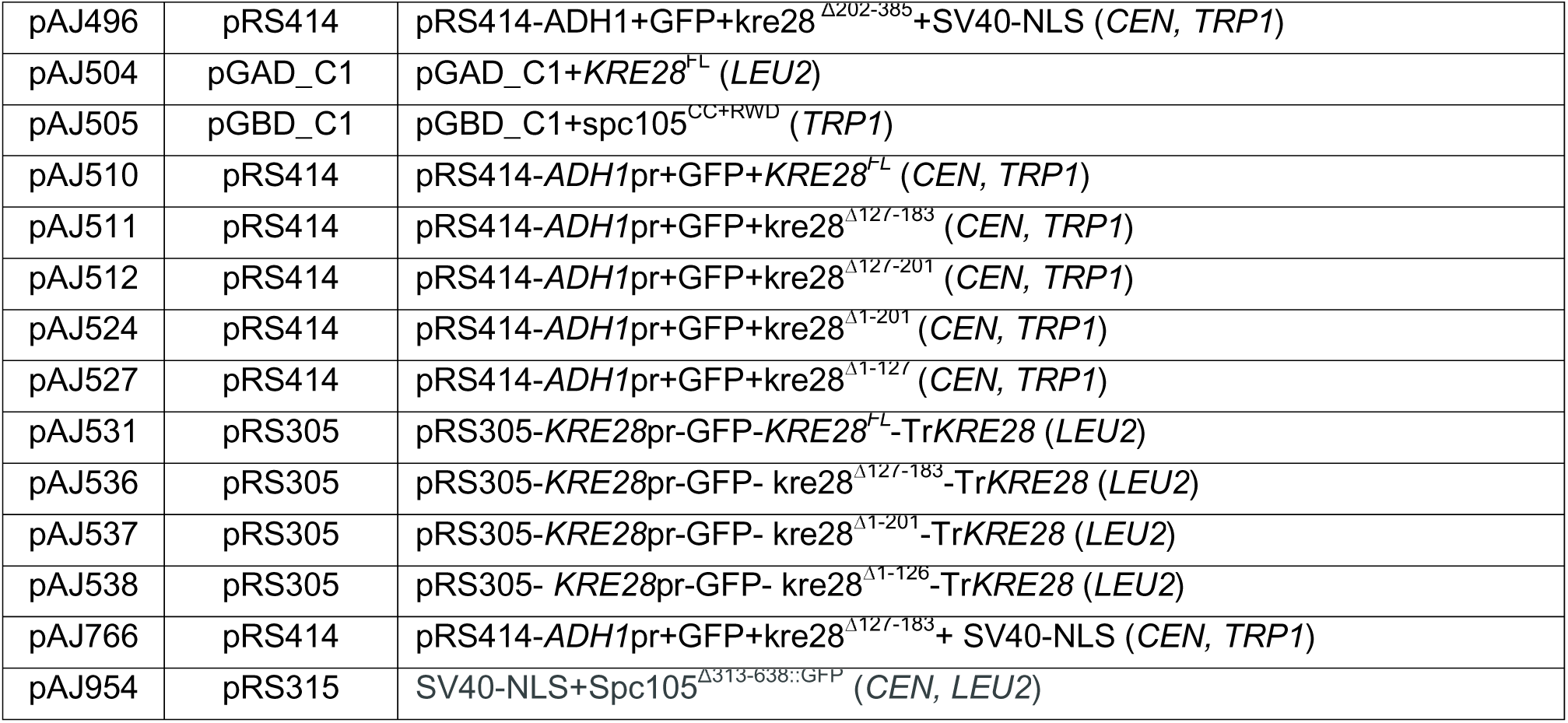
Plasmids used in this study

To construct the yeast strains with internally tagged Spc105 mentioned in Figure S1, and the strains with truncated Spc105 mentioned in Figure 3, first we used *Bst*EII digest of pRS305 chimera or *Stu*I digest of pRS306 chimera of Spc105-GFP fusion alleles to transform AJY3278 (*SPC105/*Δ*::NAT*) that was later sporulated to obtain the haploid segregants expressing only GFP fusion copy of Spc105. To construct some these strains first we deleted the genomic copy of *SPC105* in a strain already supplemented with the wild-type *SPC105* gene expressed from centromeric plasmid yCP50 (*URA3)*. Then the pRS315 chimera containing Spc105-GFP alleles were transformed in that strain. After that, the strains expressing only Spc105-GFP alleles were generated by negative selection for yCP50 on 5-FOA plates.

The construction of chimeras for overexpression of Kre28^FL^ or truncations was achieved by cloning N-terminal GFP tagged fusions of Kre28 in a centromeric plasmid pRS414, where *KRE28* ORF is flanked by *ADH1* (Alcohol Dehydrogenase 1) promoter and *GFP* ORF at the upstream and *CYC* (Cytochrome C) terminator at the downstream (obtained from Dr. Mara Duncan’s lab, department of Cell and Developmental Biology, University of Michigan). Kre28 fragments were cloned in *Bam*HI-*Pst*I sites. SV40-Nuclear localization signal (NLS) was cloned within *Pst*I-*Sal*I. For wild-type expression, Kre28^FL^ and its truncations were cloned in *Bam*HI-*Pst*I of a pRS305 plasmid. They were expressed as N-terminal GFP tagged fusions by its own promoter and terminator. *Bst*EII digests of these chimeras are transformed in AJY3298 (*kre28*Δ*::NAT*/*KRE28*-mCherry-Hyg) to check for their localization. To create diploid zygotes, two strains of *a* and α mating types are mixed with each other and spotted on YPD plate which was incubated for 3-4 hours at 30°C. To induce sporulation, diploid yeast cells were grown in YPD overnight to stationary phase. Next day cells were pelleted down and resuspended with starvation media (0.1% yeast extract, 1% potassium acetate, 0.025% dextrose) and incubated 4-5 days at RT.

### Yeast two hybrid assay

We performed yeast two hybrid experiments by co-transforming both of prey (pGAD_C1) and bait (pGBD_C1) chimera in strain AJY3802 (PJ69A) [40]. Then we streaked two of the transformants for each prey-bait pair on synthetic dextrose plates of histidine dropout (-HIS) and dropout of histidine and adenine (-HIS-ADE). Plates were incubated in 32°C for at least 3 days.

### Benomyl sensitivity assay

This experiment was performed as described previously [24, 27]. Starting from 0.1 OD_600_ of log phase cultures, we prepared 10-fold serial dilutions and frogged or spotted them on YPD and YPD containing 20 μg/ml and 30 μg/ml benomyl. At least two biological replicates were used, and spotting were repeated twice for each set of experiments. The plates were incubated at 32 °C and pictures were taken after 2 (YPD) to 3 (YPD+Benomyl) days. For space limitations, we showed only YPD+Ben_30_ plates. For all spotting assays with benomyl, we used YEF473 or strain expressing Spc105^222::GFP^ as wide-type (positive control) and *mad2*Δ *or sgo1*Δ as negative controls. Before spotting, we grew the strains expressing truncated kre28 or Kre28_FL_ control in synthetic dextrose media (Sd-Trp) without Tryptophan to culture only cells carrying the pRS414 chimera. As shown in the plate images, we also used strains expressing Kre28_FL_-mCherry along with truncated kre28 as controls which did not show any discernable difference in growth, compared to wild-type.

### Microscopy and image acquisition and analyses

A Nikon Ti-E inverted microscope with a 1.4 NA, 100X, oil-immersion objective was used for experiments mentioned in the paper [41]. A ten-plane Z-stack was acquired (200nm separation between adjacent planes). To measure Ndc80 and Spc105-mCherry, an extra 1.5x opto-var lens was used. We measured total fluorescence of kinetochore clusters (16 kinetochores in metaphase) by integrating the intensities over a 6x6 region centered on the maximum intensity pixel. We used median intensity of pixels immediately surrounding the 6x6 area to correct for background fluorescence. The calculation was performed using semi-automated MATLAB programs as described earlier [42]. FRET, high-resolution colocalization, fluorescence distribution analyses and analyses of the images were performed as previously described [6, 28, 41, 43, 44]. While measuring proximity ratio, we considered any value below 0.10 as no FRET (mean of the data marked as black), range between 0.10 and 0.3 as medium to low FRET (average of the data marked as red) and any values above 0.3 as high FRET (mean of the data marked as red).

### Titration of Kre28 and Spc105 proteins levels and quantification of Kre28, Spc105 and Ndc80 intensities

We grew the strains with *prGALL-SPC105* or *prGALL-KRE28* in presence of raffinose (2%). On the day of the assay, we supplemented the media with variable amounts of galactose as discussed previously (2%, 1.5%, 0.5%, 0.2%, 0.1% and 0.05%) [28]. We determined the number of Kre28, Spc105 and Ndc80 from their intensities as reported previously [28]. We first deduced the fluorescence intensities of Kre28-mCherry, Ndc80-GFP and Spc105-GFP from bioriented kinetochores. We used AJY939 (Ndc80-GFP, Spc25-mCherry) as a reference to obtain the intensities for known number of Ndc80 and Spc25 molecules at the bioriented kinetochores. AJY939 was cultured under same imaging conditions as the experimental strains, and the calibration data was acquired throughout the duration of this study. This calibration accounted for alteration in the microscope and imaging technique set up over time. We used the values of Ndc80-GFP and Spc25-mcherry to determine the number of molecules of Spc105-GFP and Kre28-mCherry that were loaded in the bioriented kinetochores.

### Preparation of cell lysates and western blot assay

To prepare cell lysates, log-phase cells (OD_600_ 2.0) were pelleted, resuspended in sample buffer (2% SDS, 1% 2-mercaptoethanol), boiled, and lysed by glass-bead mechanical disruption [6]. The lysates were collected after centrifugation. After separating the proteins by 10% SDS–PAGE, samples were transferred to nitrocellulose or PVDF blocked with 5% skimmed milk in TBS-T (137 mM NaCl, 15.2 mM Tris-HCl, 4.54 mM Tris, 0.896 mM Tween 20), and then probed with primary and fluorescent secondary antibodies. Mouse α-GFP antibody was from Santa Cruz Biotechnology (1:2000, GFP(B-2):sc-9996). Peroxidase conjugated α-mouse IgG (1:5000; Sigma, A-4416) treated with ECL (Thermo scientific) was used to develop the western blot.

### GFP-trap and RFP-trap assay to pull down Spc105 and immunoblot assay

We used AJY6273 and AJY6274 for GFP-trap experiments. As mentioned in the strain list, AJY6273 expresses Spc105^455::GFP^ from a centromeric plasmid (pRS315) and genomic *SPC105* allele is deleted. The truncation of 313-638 affects the cell viability, hence in AJY6374, the genomic *SPC105* allele was intact and spc105^Δ^was expressed from a centromeric plasmid (pRS315). We grew both the strains in synthetic dextrose media devoid of Leucine (SD-LEU) to maintain the centromeric plasmid before harvesting them for cell lysis. For RFP-trap assays we used AJY3483 and AJY3484 (See strain list for detailed information on their genotypes). We grew the strains in SD-TRP (Synthetic dextrose devoid of Tryptophan) media till late log-phase before harvesting the cells. We lysed the cells by glass-beads in presence of buffer H 0.15 (25 mM HEPES of pH 8.0, 2.0 mM MgCl_2_, 0.1 mM EDTA of pH 8.0, 0.5 mM EGTA-KOH of pH 8.0, 15% Glycerol, 0.1% IGEPAL-CA-630, 150 mM KCl), supplemented with 0.2 mM PMSF, protease inhibitor cocktail and phosphatase inhibitor cocktails [45]. We isolated clear lysates of the strains and from there, we incubated equal amount of lysates with pre-equilibrated beads of GFP-trap (Chromotex, gta-20) or RFP-Trap (Chromotek, rta-20) overnight. Next day, we washed the beads with post IP wash buffer (composition as mentioned above) with and without 2mM Dithiothreitol (DTT), before boiling them in presence of 1xSDS loading buffer.

After subjecting the proteins through SDS-PAGE, we transferred them to nitrocellulose membrane which we blocked with 5% skimmed milk in 1x phosphate buffered saline-Tween (PBS-T, 137 mM NaCl, 10 mM phosphate, 2.7 mM KCl, 0.05% Tween 20). We probed the blot with Mouse anti-GFP (JL8, Living Colors, Takara, 1:3000) or Mouse anti Ds-Red (Santa Cruz Biotechnology, sc-390909, 1:2000), Mouse α-Flag M2 (Sigma-Aldrich, 1:5,000) and HRP conjugated secondary anti-Mouse (Sigma aldrich, 1:10000). ECL (Thermo scientific) treatment was used to develop the western blot. We exposed the anti-Ds-Red blots to X-ray films. Anti-GFP and anti-Flag blots were imaged by C600 imager (Azure Biosystems).

We calculated the band intensities by ImageJ in RFP-TRAP assay. Then we normalized both Spc105 and Kre28 intensities of the truncation mutants with those of the full-length proteins. After that, we normalized Kre28 IP intensities with that of normalized Kre28 input values. Following that, we divided the Kre28 IP intensities with Spc105 IP intensities. We mentioned those values as fold difference at the bottom of the immunoblot panels (Figure 4D).

### Flow cytometry

We performed flow cytometry as described previously [24, 27]. For strains expressing Kre28_FL or truncated kre28, we started with overnight inoculums grown in Sd-Trp and shifted to YPD to grow till early to mid-log phase before supplementing the media with Nocodazole (final concentration 15µg/ml) or DMSO control. We collected cell samples at 0, 1, 2, 3-hour post drug treatment and fixed them with 70% ethanol before storing them at 4 C overnight. Next day, after removing the Ethanol, treated the samples with bovine pancreatic RNase (Millipore Sigma, final concentration 170 ng/ml) at 37°C for 6 h-overnight in RNase buffer (10 mM Tris pH8.0, 15 mM NaCl). After that, we removed the RNase and resuspended the cells in 1x PBS. We treated the samples with propidium iodide (Millipore Sigma, final concentration 5 mg/ml in PBS) for at least 1 h at room temperature before analyzing them using the LSR Fortessa machine (BD Biosciences) in Biomedical research core facility, University of Michigan medical school. We analyzed and organized the data using FlowJo software (FlowJo_V10.7.1_CL).

### Nuclear localization signal (NLS) mapping

We performed NLS mapping by pasting the amino acid sequences of Spc105 and Kre28 in the website of http://nls-mapper.iab.keio.ac.jp/cgi-bin/NLS_Mapper_form.cgi and keeping the cut off score to 5.0 [46, 47]. It scored SV40-NLS (TPPKKKRKVA, monopartite) as 15.5. We observed score of 9.0 (monopartite) for Spc105^337-345^ (SSNKRRKLD) and 6.9 (bipartite) for Spc105^599-625^ (NTLKREYEKLNEEVEKVNSIRGKIRKL). We did not find any NLS for Kre28 without setting the cut-off score to 4.0. Kre28^207-234^ and Kre28^286-317^ displayed NLS scores of 4.2 and 4.0 (bipartite) respectively.

### Prediction of coiled coil domains in Kre28 and spc105^451-917^

We predicted the Coiled coil domains of full-length Kre28 and spc105^451-917^ by inserting their amino acid sequences on the website of https://embnet.vital-it.ch/software/COILS_form.html [48]. We used MTIDK matrix for the prediction. We downloaded the output in postscript format and further configured by adobe illustrator.

### Prediction of spc105^451-917^- Kre28 interaction interface by Colabfold

We used Colabfold to predict the complex formed by interaction of Kre28^FL^ and _spc105_^451-917^ (https://colab.research.google.com/github/sokrypton/ColabFold/blob/main/beta/AlphaFold2_advanced.ipynb, [32]). The following parameters were used for the structure prediction: msa_method: mmseqs_2, pair_mode: unpaired, pair_cov = 50, pair_qid = 20, rank_by = pTMscore, num_models = 5, use_ptm = True, max_recycles = 3, tol = 0, num_samples = 1, subsample_msa = True, num_relax = None. We processed the figures of the predicted structure by PyMOL 2.5 (https://pymol.org/2/, Schrödinger, LLC). The pLDDT confidence matrix was depicted by spectrum b mapping in PyMOL. Pair Alignment Error matrix was prepared by Colabfold.

### Statistical analysis

We analyzed the data and assembled the graphs by GraphPad Prism 8 software. We performed unpaired t-test (Mann Whitney test) and one-way anova analyses to check the statistical significances of the data. The p-values are mentioned on the top of the graph.

## Disclosure of Potential Conflicts of Interest

There is no conflict of interest among the authors.

## Funding

This work was financially supported by R01-GM-105948 to APJ.

## Acknowledgements

We thank members of Joglekar lab for technical support, helpful discussions and positive criticism. We also like to thank Dr. Mara Duncan for providing us with reagents needed for this study and comments on this manuscript.

## Author contributions

Experiments were planned and designed: APJ BR. Performed the experiments: APJ, BR and JS. Analyzed the data: APJ BR JS. Wrote the paper: APJ BR.

## Media Summary

In our paper, we studied the localization of essential kinetochore protein Kre28 (homolog of human Zwint1) at the kinetochore microtubule attachment site (KMN network). We also report that the essential kinetochore protein Kre28 regulate the localization and organization of its binding partner Spc105 (homolog of human Knl1) in the kinetochore microtubule attachment site. This subsequently aid in proper maintenance of the spindle assembly checkpoint signaling.

## Supporting information

**Supporting figure S1: Related to figure 1.**
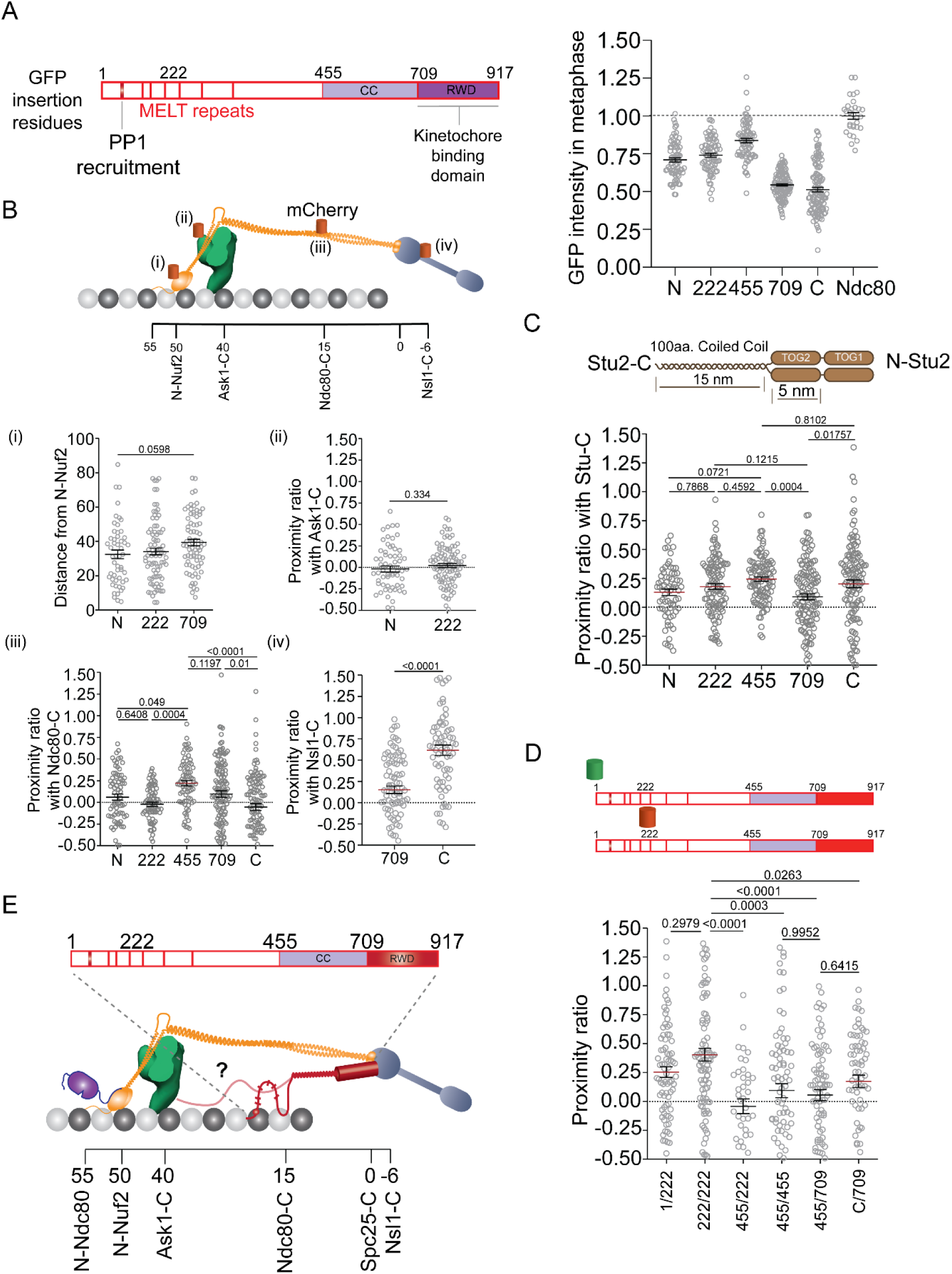
**The N terminal phosphodomain of Spc105 (1-455) is highly disordered in nature.** (**A**) Left: Line diagram of Spc105 molecule. The illustration was reproduced from our previous study [24]. Red bars represent PP1/Glc7 recruitment site (amino acid 75-79), and six MELT repeats. Amino acid locations of GFP fusion are shown at the top on N (1), amino acid 222, 455, 709 and C (917). Right: Scatter plot shows intensities of Spc105 (GFP at N, 222^nd^, 455^th^, 709^th^ amino acids and at C termini) in bioriented kinetochores of yeast. Intensity of Ndc80-C is mentioned as a control. (**B**) Organization of Spc105 with respect to Ndc80, Mtw1 and Dam1 complexes. Top: The organization of kinetochore proteins in a metaphase kinetochore of yeast along the microtubule axis. Positions of C termini of Nsl1 (-6nm), Spc25/Spc24 (0nm), Ndc80/Nuf2 (15nm) and Dad4/Ask1 (40nm) and N termini of Nuf2 (50nm) and Ndc80 (55nm) are shown. Red/orange barrel are shown to indicate mCherry tagging of N-Nuf2, Ask1-C, Ndc80-C and Nsl1-C in individual yeast strain. Bottom: (i) Scatter plot displaying distance between the centroids of 222, 455 and 709^th^ amino acid positions of Spc105 and GFP-Nuf2 (N-Nuf2). At least 49 kinetochore foci were analyzed to obtain this data. Scatter plot of proximity ratio [directly proportional to FRET (Förster resonance energy transfer) [6]] with mean± s.e.m showing proximities of different amino acid positions of Spc105 molecules (222 for Spc105^222::GFP^, 455 for Spc105^455::GFP^, 709 for Spc105^709::GFP^, and C of Spc105-C) with Ask1-C (ii), Ndc80-C (iii) and Nsl1-C (iv). Minimum number of kinetochore foci analyzed to acquire this data: 62 for Ask1-C, 75 for Ndc80-C and 76 for Nsl1-C. The p-values obtained by t-test or one-way anova test are mentioned above the graphs. We observed a high FRET between Nsl1-C and Spc105-C but the FRET decreases if the donor is moved to Spc105 709^th^ amino acid. We noticed a mid to low FRET between Ndc80-C and Spc105^455^. (**C**) Organization of Spc105 with respect to Stu2, marker protein of microtubule plus ends. Top: Schematic diagram of Stu2 dimer [41, 49]. Bottom: Scatter plot with mean±s.e.m showing proximity ratio of different amino acid positions of Spc105 with Stu2-C. At least 105 kinetochore foci were analyzed for this plot. The p-values obtained from one-way Anova test are mentioned at the top of plot. The position of MT plus tip and hence the position of Stu2 may vary from one kinetochore to another [6]. Our observations indicated low to moderate proximity for all the GFP insertion points of Spc105 that suggests the whole molecule of Spc105 remains in close proximity with MT lattice. (**D**) Proximity ratio showing FRET among the adjacent Spc105 molecules in metaphase kinetochores. Minimum number of kinetochore foci examined for this graph is 40. Our observations are consistent with the previous studies which hypothesized that the Spc105 phosphodomains remain in close proximity with each other to create interaction foci of SAC proteins in unattached KTs [14, 50]. (**E**) Localization of Spc105 molecules with respect to other proteins in the KMN network of bioriented kinetochores. The illustration was reproduced from our previous study [24]. Although the C-terminus of this protein (amino acid 709-917) is structured and could be localized in the proximity of C-termini of Spc24/Spc25 and Nsl1, the N-terminal region (amino acid 1-455) is unstructured and could be localized within a 20nm region between C-termini of Dad4 and that of Ndc80.

**Supporting figure S2: Related to figure 1, 2 and 4.**
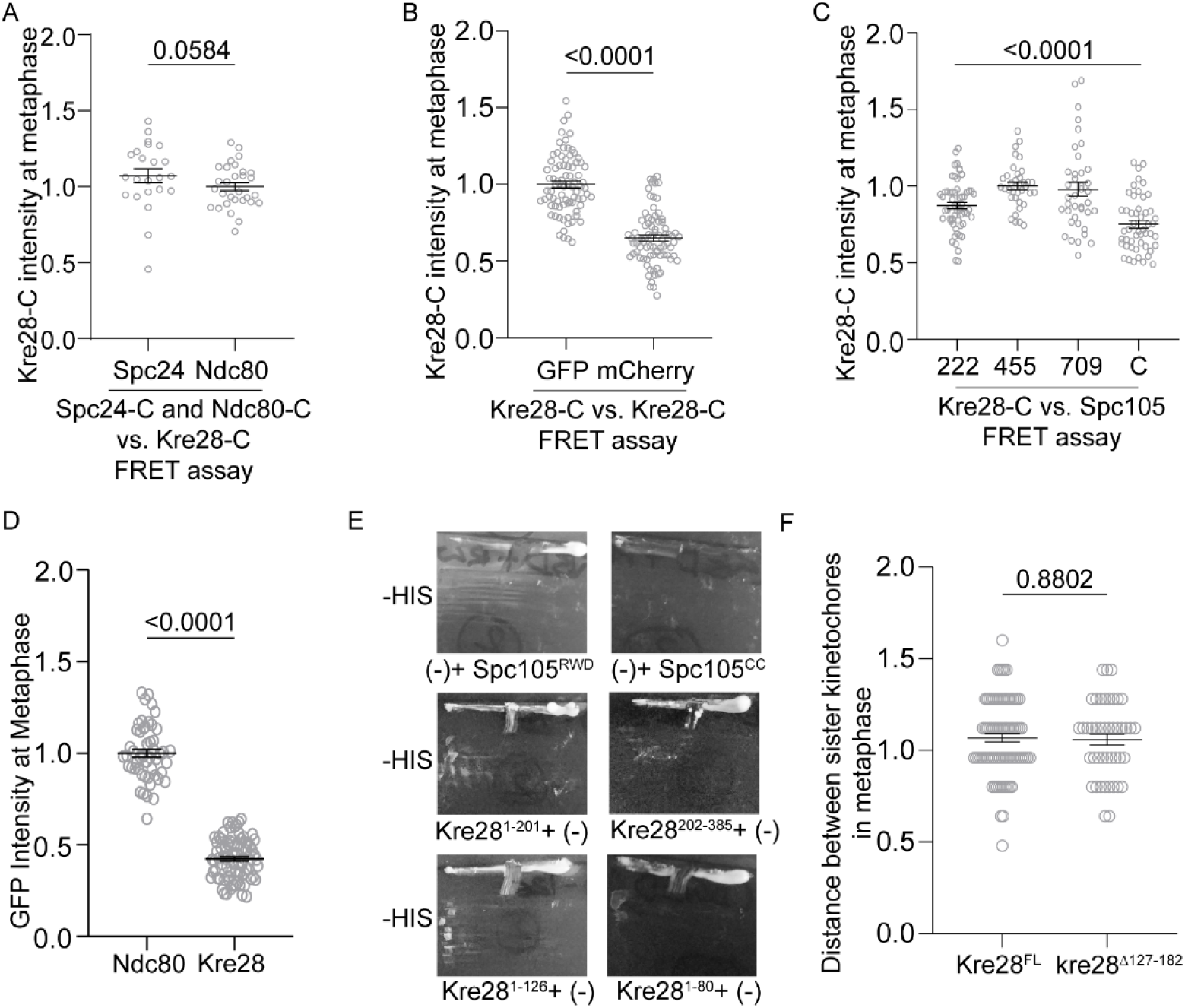
(**A-C**) Normalized intensities of Kre28-C in 3 sets of FRET experiments. The dot plots are showing mean±s.e.m of normalized intensities of Kre28-GFP or Kre28-mCherry. Data in (**A**) was normalized by Kre28 intensity in FRET assay with Ndc80-GFP. Data in (**B**) was normalized by Kre28- GFP. Data in (**C**) was normalized was Kre28 intensity in FRET experiment involving Spc105^455::GFP^. (**D**) Normalized GFP intensities of Ndc80 and Kre28, localized at bioriented kinetochores. (**E**) The plate images show control experiments of yeast two hybrid assay performed in Figure 2 A. (**F**) Dot plot displays mean+s.e.m of distances between sister kinetochores in metaphase cells expressing Kre28^FL^or kre28^Δ127-183^. At the top of the plot, the p-value that was derived from the unpaired t-test done on the data, is mentioned.

**Supporting Figure S3, related to Figure 2 and 3.**
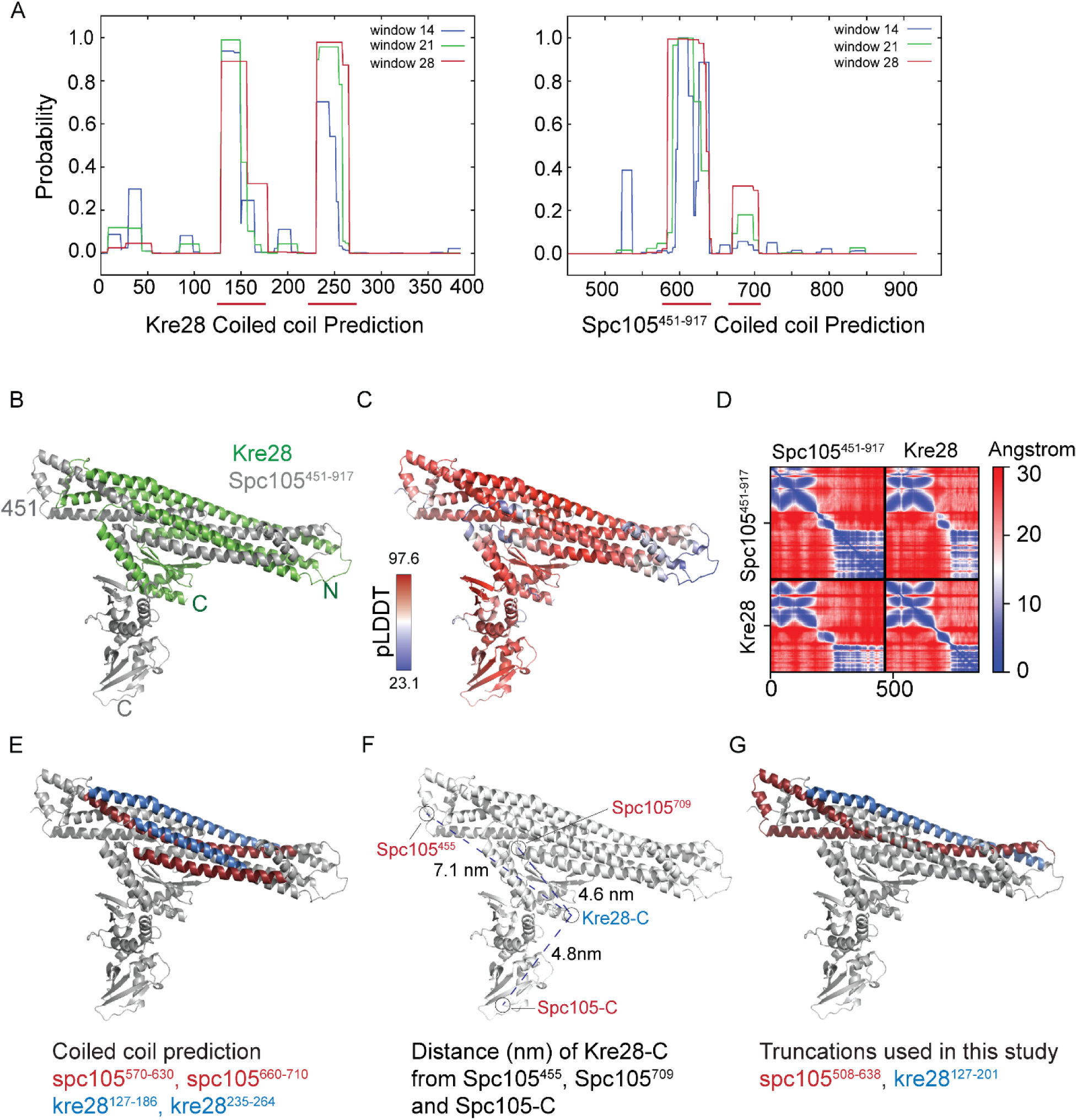
Prediction of coiled coil domains of Kre28 and that of spc105^451-917^ and their binding interface: (A) Prediction of coiled coil domains in Kre28^FL^ and spc105^451-917^ amino acid sequences by COILS server (https://embnet.vital-it.ch/software/ COILS_form.html). Red lines at the bottom of the plots depicts the domains of Kre28 (approximately kre28^125-175^ and kre28^235-265^) and spc105^451-917^ (approximately spc105^570-630^ and spc105^660-710^) which showed high scores in COILS prediction. (**B**) Predicted backbone structure with the highest pTMscore for the 1:1 heterodimer of Kre28-Spc105^451-917^ using Colabfold. Kre28 and spc105 are shown in green and gray colors respectively. Amino (spc105^451^) and Carboxy termini (C) are also indicated for both the proteins. The following parameters were used for the structure prediction: msa_method: mmseqs_2, pair_mode: unpaired, pair_cov = 50, pair_qid = 20, rank_by = pTMscore, num_models = 5, use_ptm = True, max_recycles = 3, tol = 0, num_samples = 1, subsample_msa = True, num_relax = None. Structure visualizations were generated by Pymol. (**C**) Structure of the heterodimer backbone colored by the pLDDT confidence matrix. The color bar shows the degree of confidence with high confidence in red and low confidence in blue. (**D**) Pair Alignment Error matrix for the displayed structure. (**E**) Mapping of the domains of Kre28 (kre28^125-175^ and kre28^235-265^, colored in blue) and that of spc105^451-917^ (spc105^570-630^ and spc105^660-710^, colored in red) on the predicted heterodimer, which have high scores in the coiled coil prediction (see Figure S3A). (**F**) Distances of alpha carbon atom of Kre28^385^ (Kre28-C) from that of Spc105^455^ (7.1 nm), Spc105^709^ (4.6 nm) and Spc105^917^ (Spc105-C) (4.8 nm) in the predicted structure (shown by blue dashed line). Black circles depict the positions of residues on the structure. (**G**) Plotting of the domains of Kre28 (kre28^127-201^, colored in blue) and that of spc105^451-917^ (spc105^507-638^, colored in red) on the structure, that were truncated in our study.

## Notes

### Competing Interest Statement

The authors have declared no competing interest.

